# Immuno-functionomics reveals geographical variation and a role for TLR8 in mRNA vaccine responses

**DOI:** 10.1101/2025.03.28.645987

**Authors:** W. Huisman, S. Azimi, Y.N. Nguyen, A.C. de Kroon, K. de Ruiter, D.L. Tahapary, M.D. Manurung, C.R. Pothast, Y.C.M. Kruize, M.H.M. Heemskerk, T. Supali, L.G. Visser, A.H.E. Roukens, M. Yazdanbakhsh, S.P. Jochems

## Abstract

The innate immune system plays a pivotal role in pathogen defense via pattern recognition receptor sensing, initiating responses upon infection or vaccination. Understanding its functional capacity is crucial for deciphering correlates of vaccine efficacy and understanding responses to infection. In this study, we developed a holistic approach to study immune function, generating >3100 readouts across 16 cell types, 18 pattern recognition receptors and 11 produced cytokines using spectral flow cytometry. To explore geographical variation, we studied the immune system of Europeans, urban, and rural Indonesians. We found differences in immune responses, such as increased IL1β production in rural Indonesians and impaired IFNγ production by innate lymphocytes after TLR8 stimulation. In Europeans vaccinated with mRNA-1273, baseline IFNγ production by innate lymphocytes correlated with SARS-CoV-2 Spike-specific immune responses. *In vitro* mRNA vaccine stimulation also induced IFNγ production, which was TLR8-dependent and reduced in rural Indonesians. This study highlights functional immune diversity and TLR8’s potential role in mRNA vaccine responses.

**Summary:** We developed an approach to study the function capacity of the immune system, exploring geographical variation and mRNA vaccine responses. This revealed TLR8’s potential role in responding to mRNA vaccines, and an impairment in this pathway in rural Indonesians.

## Introduction

The innate immune system plays a critical role in protecting against infections and shaping adaptive immunity. It consists of various cell types that express unique combinations of pattern recognition receptors (PRRs) which sense pathogen-associated molecular patterns (PAMPs)^1^. These PRRs can be classified based on their location and type of PAMPs they detect. For instance, extracellular PRRs, such as Toll-Like Receptors (TLR) 1, 2, 4, 5, 6 and Dectin1 and 2 recognize components from bacteria, parasites, and fungi. Meanwhile, endosomal and cytosolic PRRs, like TLR3, 7, 8, 9, RIG-I, STING, and cGAS, detect nucleic acids and are essential for sensing intracellular pathogens, including viruses. When activated, PRRs initiate signalling cascades involving nuclear factor-kappa B (NF-kB) and/or interferon regulatory factors (IRFs) leading to the production of interferons and proinflammatory cytokines, which in turn shape adaptive immunity^2^.

The importance of reactivity of the innate immune system towards vaccines has already been shown before, whereby levels of activated innate cells *in vivo* early after vaccination consistently correlate with the induction of robust adaptive immunity^3, 4^. Systems biology approaches have identified potential predictive gene signatures of good universal vaccine responses^5-9^, like enriched proinflammatory response genes downstream of NF-kB^9^, and/or T-cell enriched modules and B-cell receptor signalling^5^. However, these signatures were mainly predictive for vaccine responses in young adults and could not be validated in cohorts of older adults, nor did they include a diverse geographical variation. Indeed, predicting vaccine responses based on baseline phenotypic immunological data remains challenging, as cellular phenotype and transcriptome do not necessarily reflect functional capacity.

The individual and population-level variability in immune responses is significant, with factors contributing to this variation that are not yet fully understood. Notably, vaccine immunogenicity is often suboptimal in populations most at risk from infectious diseases, including older adults, infants, and individuals in low- and middle-income countries (LMIC). Various factors may contribute to the reduced vaccine immunogenicity observed in individuals from LMIC^10^, such as chronic exposure to environmental pathogens, parasites, and co-infections, which can lead to an anti-inflammatory or hyporesponsive state of the immune system. This affects not only the immunogenicity of vaccines, but also modulates responses to infections, such as SARS-CoV-2^11^. Even within a country the state of the immune system can vary^12^, as people in rural areas of LMICs often have greater exposure to microbes and parasites than people living in urban areas^13, 14^. This is mirrored in the immunological phenotype, as for example by comparing the immune system of Indonesians living on rural Flores island to Indonesians living in the Indonesian capital of Jakarta, it was found that the immune system of urban Indonesians was more similar to the immune system of Europeans^12^. However, despite the described general hyporesponsiveness and observed differences in trained immunity^15^, the exact pathways of the innate immune system affected remain unclear.

Our study aimed to explore the functional capacity to different PRR stimulations of the innate immune system using an ‘Immuno-functionomics’ approach. We hypothesized that functional responses would vary in a cell-type and PRR-specific manner across individuals from different geographic regions and could predict vaccine responses. We found that activation of innate lymphocyte populations following TLR8 stimulation was reduced in rural Indonesians. Activation of these innate lymphocyte populations are also predictive of mRNA vaccine responses in an independent European vaccination cohort. *In vitro* responses to mRNA stimulation were reduced in rural Indonesians, and could be abrogated by blocking TLR8. This highlights the potential for predicting vaccine responses based on functional capacity, allows identification of differences in capacity across regions and revealed a previously unappreciated role for TLR8 sensing in mRNA vaccine responses.

## Material and methods

### Collection of donor material and sample selection

All participants provided written informed consent before sampling according to the Declaration of Helsinki. Rural Indonesian samples were obtained as part of the SugarSPIN trial (ISRCTN; doi.org/10.1186/ISRCTN75636394), a household-based cluster-randomized double-blind trial on investigating the association between whole-body insulin sensitivity and helminth infections, that was conducted in three rural villages in Nangapanda, Ende district of Flores Island (East Nusa Tenggara), Indonesia^16^. This study has been approved by the ethical committee of Faculty of Medicine Universitas Indonesia (ref: 549/H2.F1/ETIK/2013). For this analysis, baseline samples prior to any potential albendazole treatment were included.

Age- and sex-matched healthy volunteer samples, consisting of Caucasians from the Netherlands and individuals from urban centres in Indonesia, such as central Jakarta, were obtained from the LUMC Blood Bank (L19.002) or the LUMC Biobank for Infectious Diseases and included in this study. We also included samples from Dutch recipients of Moderna mRNA-1273 vaccine who participated in a study where the immunogenicity of the intradermal route administration was compared to intramuscular injection^17^ (IDSCOVA trial; EUCTR2021-000454-26-NL). In this trial, healthy adults between 18 and 30 years without laboratory-confirmed or self-reported SARS-CoV-2 infection were recruited. Only samples from SARS-CoV-2 naïve individuals without SARS-CoV-2 specific antibodies, T and B cells at baseline were included in this analysis. For all individuals (**Table 1**), PBMCs were isolated by standard Ficoll-Isopaque separation and stored in the vapor phase of liquid nitrogen.

**Table 1.**
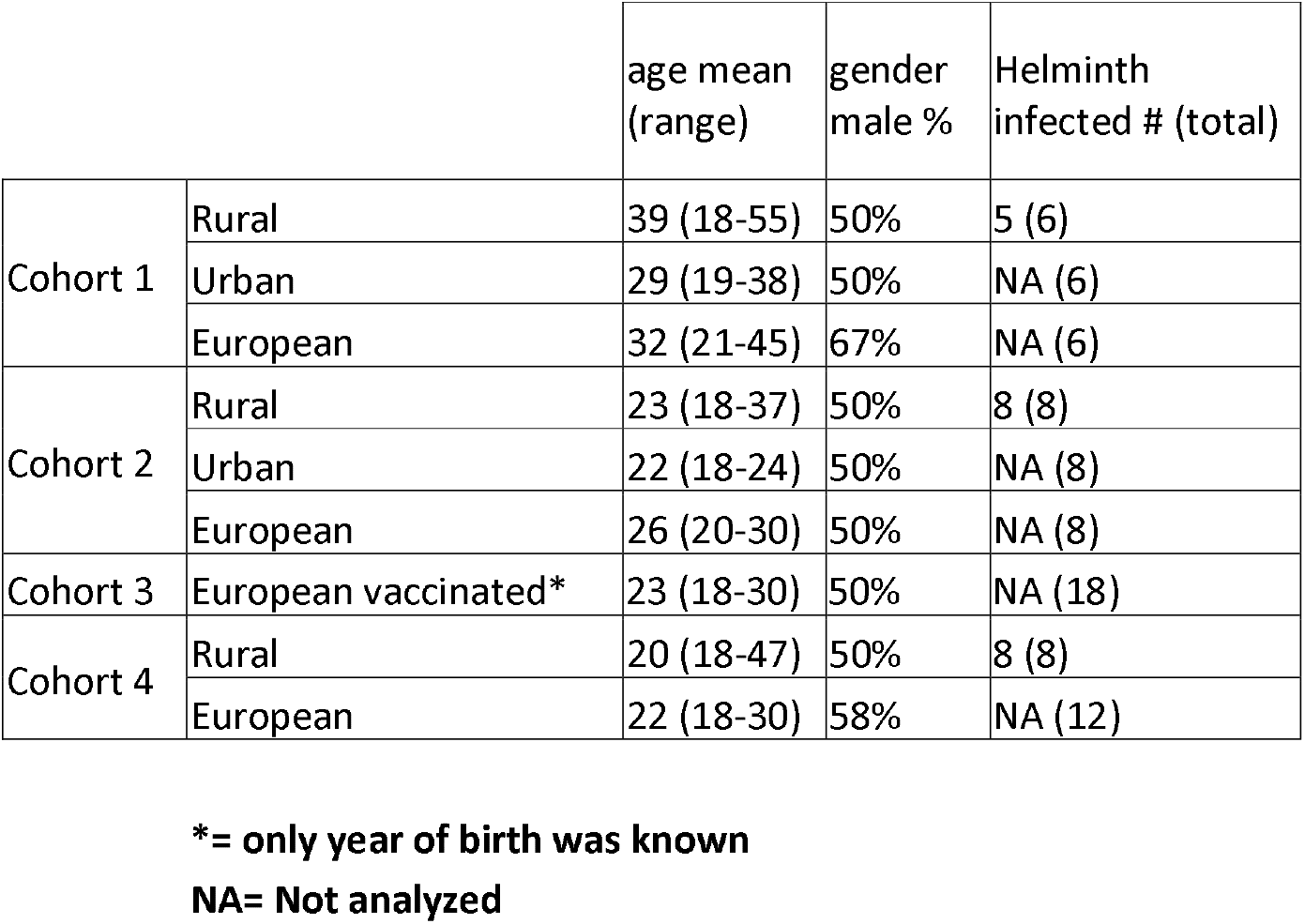
General participant characteristics.

|  |  | age mean<br>(range) | gender<br>male % | Helminth<br>infected # (total) |
| --- | --- | --- | --- | --- |
| Cohort 1 | Rural | 39 (18-55) | 50% | 5 (6) |
|  | Urban | 29 (19-38) | 50% | NA (6) |
|  | European | 32 (21-45) | 67% | NA (6) |
| Cohort 2 | Rural | 23 (18-37) | 50% | 8 (8) |
|  | Urban | 22 (18-24) | 50% | NA (8) |
|  | European | 26 (20-30) | 50% | NA (8) |
| Cohort 3 | European vaccinated* | 23 (18-30) | 50% | NA (18) |
| Cohort 4 | Rural | 20 (18-47) | 50% | 8 (8) |
|  | European | 22 (18-30) | 58% | NA (12) |

**\*= only year of birth was known**
**NA= Not analyzed**

### *In vitro* Immuno-functionomics stimulations using 18 different PRR-agonists

PBMCs were thawed in a 37°C water bath and transferred to a 15mL centrifuge tube (Greiner, Germany) and washed by slowly adding 1 mL pre-warmed thawing medium (Roswell Park Memorial Institute (RPMI) 1640 Medium (Gibco, United Kingdom), 20% heat-inactivated fetal bovine serum (FBS) (PAN Biotech, United Kingdom), 100 µg/mL streptomycin (Gibco, United Kingdom) and 100 U/mL penicillin (Gibco, United Kingdom) (Pen/Strep). Another 4 mL of pre-warmed thawing medium was slowly added to the cell suspension. After cells were spun down 5 minutes at 450g, another 5 mL of pre-warmed thawing medium was added and cells were resuspended. After another centrifugation and removal of supernatant, cells were resuspended in RPMI medium supplemented with 10% FBS, Pen/Strep and 2.7 mM L-Glutamine (Merck, Germany) (culture medium) at 5 million cells/mL. Then, 500.000 cells (100ul in culture medium) were plated per well in 2 round bottom 96 well Nunclon™ Delta Surface plates (Thermo Fisher Scientific, Denmark). Containing in total up to 18-PRR agonists and 2x medium controls that were stimulated either overnight or for 4 hours (**Table 2**). For the overnight stimulation plate, cells were first rested 2-4 hours at 37°C, 5% CO_2_, before 100ul of 10 PRR-agonists or medium control was added. After overnight stimulation (approximately 16 hours), 10 µg/mL Brefeldin A (Invitrogen, USA) was added, and incubated for another 4 hours. For the 4 hour stimulation plate, cells were rested overnight at 37°C, 5% CO_2_ and the following day 8 PRR-agonists or medium control was added and incubated for 1 hour. Brefeldin A was then added and incubated for another 3 hours at 37°C, 5% CO_2_. If less than 10 million cells were recovered from a thawed vial, the stimulations to use were prioritized in accordance with the order of Table 2.

**Table 2.** Pattern recognition receptor agonists used to investigate the function of the innate immune system.

| # | PRR | Agonist | stimulation:<br>4 hours or<br>O/N | company | Catalogue | Working<br>concentration |
| --- | --- | --- | --- | --- | --- | --- |
| 1 | TLR1/2 | PAM | O/N | InvivoGen | tlrl-kit1hw | 1µg/ml |
| 2 | TLR3 | Poly(I:C) | O/N | InvivoGen | tlrl-kit1hw | 10µg/ml |
| 3 | TLR4 | LPS | 4 hours | InvivoGen | tlrl-kit1hw | 5µg/ml |
| 4 | TLR5 | FLAST | 4 hours | InvivoGen | tlrl-kit1hw | 1µg/ml |
| 5 | TLR7 | Imiquimod | 4 hours | InvivoGen | tlrl-kit1hw | 2µg/ml |
| 6a | TLR9 | ODN2006 | 4 hours | InvivoGen | tlrl-2006 | 15µg/ml (2µM) |
| 6b | TLR9 | ODN2395 | 4 hours | InvivoGen | tlrl-2395 | 15µg/ml (2µM) |
| 7 | Dectin 2 | Furfurman | O/N | InvivoGen | tlrl-ffm | 1µg/ml |
| 8 | TCR-MHCI/II | SEB | 4 hours | Sigma-Aldrich | S4881 | 200ng/ml |
| 9a | IL-12R | IL-12 | O/N | InvivoGen | rcyc-hil12 | 50ng/ml |
| 9b | IL-18R | IL-18 | O/N | InvivoGen | rcyec-hil18 | 100ng/ml |
| 10a | TSLP-R | TSLP | O/N | Biolegend | 582406 | 500ng/ml |
| 10b | ST2 | IL-33 | O/N | Biolegend | 581806 | 500ng/ml |
| 10c | IL-17R | IL-25 (IL-17E) | O/N | Biolegend | 598902 | 500ng/ml |
| 11 | TLR8 | ssRNA40 | O/N | InvivoGen | tlrl-kit1hw | 2µg/ml |
| 12 | NOD2 | L18 MDP | 4 hours | InvivoGen | tlrl-lmdp | 10µg/ml |
| 13 | TLR2 | HKLM | 4 hours | InvivoGen | tlrl-kit1hw | 10E8 cells/ml |
| 14 | STING | 2',3'cGAMP | 4 hours | InvivoGen | tlrl-nacga23-02 | 10µg/ml |
| 15 | RIG-1 | 3p hpRNA * | O/N | InvivoGen | tlrl-hprna | 3µg/ml |
| 16 | cGAS | poly(dG:dC)* | O/N | InvivoGen | tlrl-pgcc | 0.5µg/ml |
| 17 | NOD1 | C12 iE DAP | O/N | InvivoGen | tlrl-c12dap | 1µg/ml |
| 18 | Dectin 1 | β Glucan | 4 hours | InvivoGen | tlrl-bgp | 10µg/ml |
| * |  | LycoVec <sup>TM</sup> |  | InvivoGen | lyec-1 |  |

**Abbreviations; PRR, pattern recognition receptor; O/N, overnight; TLR, Toll-like receptor**

### Flow cytometry staining

Following incubation with Brefeldin A, PBMCs were detached from the wells using 2 mM UltraPure™ EDTA, pH 8.0 (Invitrogen, USA) for 10 minutes at room temperature (RT). Cells were pelleted for 5 minutes, at 450xg at RT. For each individual stimulation, PBMCs were barcoded with 50 µL of a combination of self-conjugated CD45-CF568 (Biolegend, USA. Cat.368502) using Mix-n-Stain™ CF® Dye Antibody Labeling Kit (Biotium, Cat.92235), CD45-PacificBlue (Biolegend, Cat.368540) and/or CD45-AlexaFluor700 (Biolegend, Cat.368514) for 15 minutes at RT, followed by washing with FACS buffer (Phosphate buffered saline (PBS) containing 2 % (v/v) BSA, 2 mM EDTA) (**Supplementary Figure 1A**). Cells were washed with sterile PBS (Fresenius Kabi, Germany) twice and pooled per plate in a 15 mL falcon tube. After centrifuging and removing supernatant, cells were resuspended and stained in 50 µl for 15 min at RT with Live/Dead Blue fixable Blue Dead Cell Stain (Invitrogen, USA) solution at 1:500 dilution mixed with anti-human Fc Receptor (FcR) Binding Inhibitor (Invitrogen, USA) and True-Stain Monocyte Blocker (Biolegend) at 1:50 dilution. Then extracellular staining was performed by adding 50 µl of 2x concentrated mix containing 16 monoclonal antibodies directed against extracellular proteins **(Table 3; #1-#16)** prepared in FACS buffer with 5 µl BD Horizon™ Brilliant Stain Buffer Plus (BD Biosciences, USA) for 15 min at RT. After two washes with 80 µl and 180 µl of FACS buffer, cells were fixed with 180 µl of 4% final concentration formaldehyde solution (w/v), methanol free (Thermo Scientific, USA) for 20 minutes at 4°C. Before staining with an intracellular antibody cocktail, cells were washed with 180 µl 1xFACS buffer and fixed cells were centrifuged at 800g for all subsequent steps. Cells were permeabilized in 180 µl of 1x BD Perm/Wash buffer (BD Biosciences, USA) for 15 minutes at RT and then pelleted by centrifugation. Finally, intracellular staining was performed in 100 µl for 30 minutes at 4°C with a mix containing 13 different monoclonal antibodies directed against cytokines/interleukins prepared in 1x BD Perm/Wash buffer with 10 µl BD Horizon™ Brilliant Stain Buffer Plus **(Table 3; #17-#29)**. Monoclonal antibodies directed against Th2 cytokines IL4, IL5 and IL13 were all conjugated with PE and were measured as 1 output (Th2 cytokines). After intracellular staining, cells were washed twice with Perm/Wash buffer and finally resuspended in 400 µl FACS buffer for acquisition. Cells were acquired with fluidics boost on a 5-laser Aurora Cytometer (Cytek Biosciences, Inc, USA). Reference controls were made using either AbC™ Total Antibody Compensation Beads (Invitrogen, USA) or PBMCs **(Table 3)**. An additional unstained control, gated on monocytes (FSC/SSC) was used for unmixing.

**Table 3.**
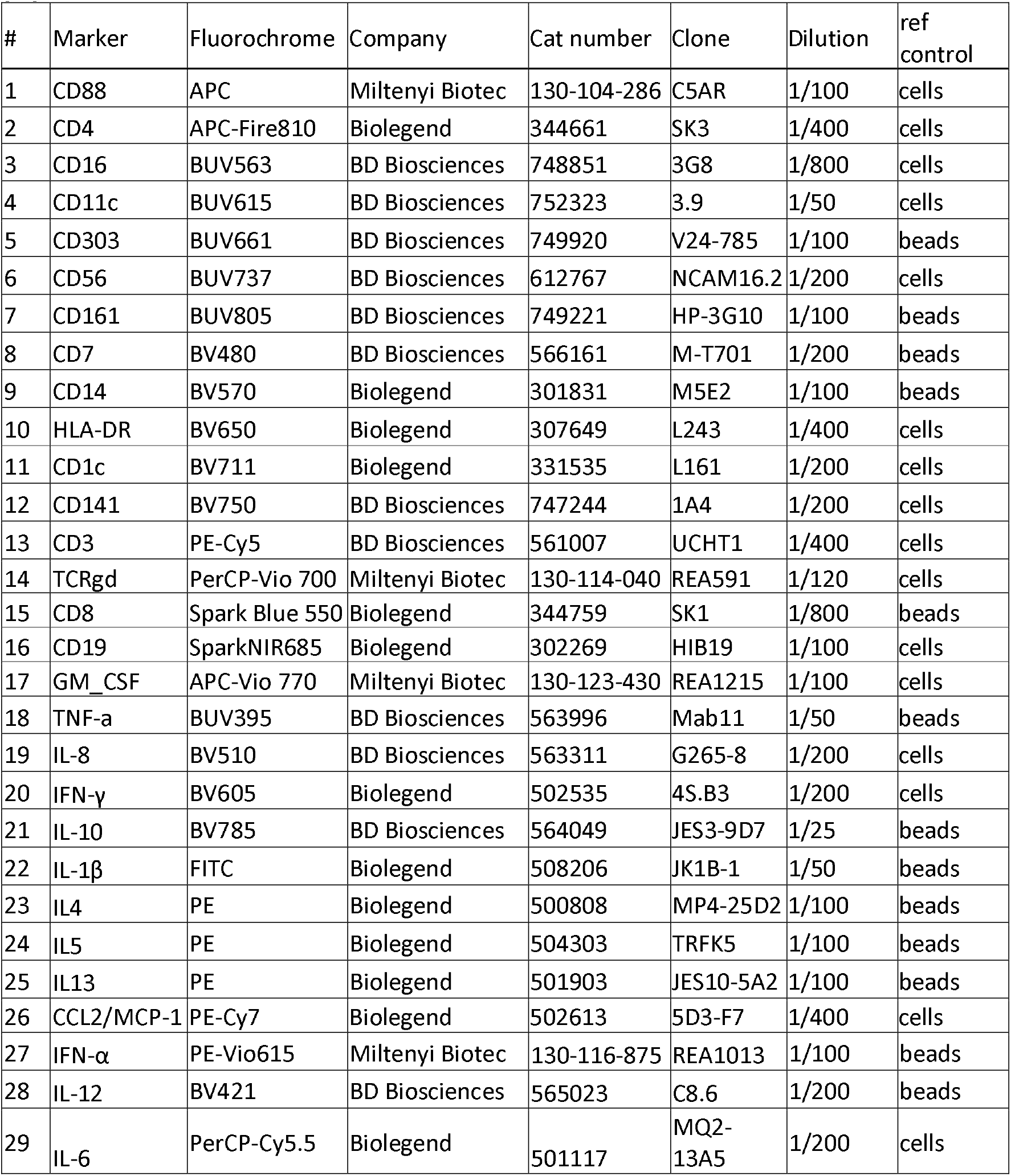
Panel of different monoclonal antibodies used for analysis of the function of immune cell populations.

For expression of different TLRs, 1 million PMBCs of individuals from cohort 2 were thawed similarly as before. Cells were resuspended and stained in 50 µl for 15 min at RT with Live/Dead Blue fixable Blue Dead Cell Stain (Invitrogen, USA) solution at 1:500 dilution mixed with anti-human Fc Receptor (FcR) Binding Inhibitor (Invitrogen, USA) and True-Stain Monocyte Blocker (Biolegend) at 1:50 dilution. Extracellular staining **(Supplementary Table 1; #1-#16)** and intracellular staining **(Supplementary Table 1; #17-#20)** were then similarly conducted as described above. Cells were acquired with fluidics boost on a 5-laser Aurora Cytometer (Cytek). Reference controls were made using either AbC™ Total Antibody Compensation Beads (Invitrogen, USA) or PBMCs **(Table 4)**. An additional unstained control, gated on monocytes (FSC/SSC) was used for unmixing.

### Flow cytometry gating and analysis

After acquisition, stimulations were manually debarcoded using FlowJo software (BD) and conditions were exported as separate .fcs files according to the barcoding scheme **(Supplementary Figure 1A and 1B)**. Additionally, immune cell populations (13 innate and 3 adaptive) were manually gated and exported as .fcs files using the gating shown in **Supplementary Figure 2**. The cDC1 subset was not exported due to the low number of cells that could be analyzed. The exported .fcs files of the different barcodes and populations were then combined in R resulting in 320 (16 populations x 20 stimuli) conditions for each sample. Cytokine levels were arcsinh transformed prior to gating. TNFα, CCL2 and IL1β were transformed with co-factor 1500 while IL12, IFNγ, IL10, IL6, IFNα and GM-CSF were transformed with co-factor 5000. IL8 and Th2 cytokines were transformed with cofactor 15.000. Cytokines were then gated in R per samples using a semi-automated process to obtain frequencies of cytokine producing cells for each condition and cell type. For each sample, cytokine and population, the 95% quantile in the medium control of the overnight or 4-hour stimulation was determined. This value multiplied by a factor 2, 2.5 (IL12 and type 2 cytokines) or 2.7 (IL6) to set a cutoff above which cells were defined as being cytokine positive. For IL8 and IL12, an upper arcsinh transformed limit of cut-off of 2 was further defined, and for IFNα the threshold was manually set at 2 regardless of the population. As IL1β and CCL2 were spontaneously produced by certain immune populations, the threshold for these cytokines was defined using the CD4^+^ T cell population and applied to all populations. Thresholds and producing cytokines were visualized for all samples and combinations of cytokines, populations and stimulations to verify correct thresholding. All R scripts used for combining exported .fcs files, transformation and semi-automated gating are available on github.

### *In vitro* stimulations with Moderna mRNA-1273 and Comirnaty Omicron mRNA-XBB 1.5

Stimulations with mRNA-1273 (Moderna, USA) and Comirnaty Omicron mRNA-XBB 1.5 (BioNTech/Pfizer, USA) were similarly conducted as for the 10 PRR-agonists that were used overnight. The mRNA-1273 vaccine was dissolved in 500 µl of the aqueous buffer that comes with the vaccine (200ug/ml). The Comirnaty Omicron mRNA-XBB 1.5, was pre-dissolved in 3000 µl of the aqueous buffer (100 µg/ml). Stimulations were performed by adding 100 µl of the dissolved mRNA-1273 or mRNA-XBB to the wells from the 96 well plate containing cells. For stimulations where the TLR8 inhibitor CU-CPT9a (Invivogen, USA) was added, cells were prior to stimulation pre-incubated with 10 µM of CU-CPT9a for 3 hours at 37°C. Vaccines and controls were added without washing, resulting in a 5 µM CU-CPT9a concentration during the overnight stimulation. Stimulations were then barcoded similarly as before using CD45-PacificBlue (Biolegend, Cat.368540), CD45-Realblue 780 (BD, Cat. 568747), CD45-Pe-cy7 (BD, Cat. 557748) and/or CD45-AlexaFluor700 (Biolegend, Cat.368514) for 15 minutes at RT.

### Statistics

Results were analyzed in R version 4.32 and Prism Version 10 (GraphPad, San Diego, CA). Statistical differences between rural Indonesians, urban Indonesians and Europeans were compared using the Limma package (v3.54.2) for all conditions that were present in at least 3 individuals per group with an average frequency >0.5% **(Supplementary Figure 3)**. Correlations between the baseline functional responses and adaptive immunity following mRNA vaccination were calculated using spearman correlation (Hmisc package v.4.8.0). Bonferroni corrections were used for the comparison of multiple groups. Heatmaps were created with the pheatmap package (v1.0.12).

## Results

### Simultaneous characterization of innate and adaptive immune cell functionality

In this study, we measured the production of 11 different cytokines by 16 immune cell populations in response to stimulation with 18 different stimuli, generating approximately 3100 functional readouts per sample **(Figure 1A, 1B and Supplementary Figure 2)**. Of these combinations, 462 showed a cytokine production of at least 0.5% relative to the parent cell population **(Figure 1B and Supplementary Figure 3)**. All populations produced multiple cytokines in response to different PRR agonists, except for B cells and CD16^+^ NK cells **(Figure 1B)**. B cells only produced TNFα in response to TLR9 stimulation, while CD16^+^ NK cells only produced IFNγ in response to stimulation with recombinant IL12 and IL18, or after TLR8 stimulation **(Figure 2A)**. CD16^-^ and CD56^Hi^ NK cells produced IFNγ, TNFα and GM-CSF in response to several ligands, consistent with their known increased ability to produce cytokines compared to the CD16^+^ NK cell population^18^ **(Figure 1B and Figure 2B)**. In line with literature, responses to some PRR-agonists were more predominant for specific populations than others. For example, TLR7 is known to be mainly expressed by pDCs^19^ and these cells produced large amounts of TNFα and IFNα in response to TLR7 stimulation **(Figure 2C)**. Contrary to TLR7, TLR8 is more broadly expressed by different monocyte and DC subsets^20^ and they produced IL1β, CCL2, TNFα, IL6, IL8 and IL12 in response to TLR8 **(Figure 2D)**. Although innate lymphoid populations (e.g. NK subsets, ILCs, DN T and DP T) do not express TLR8, they produced IFNγ in response to TLR8 stimulation **(Figure 1B and Figure 2B)**, which has been shown to be due to accessory cytokine (IL12 and IL18) release by myeloid populations^21-23^. Moreover, as expected, the type of produced cytokines also varied between immune populations. For example, cDC2s were among the largest IL12 producers, while type 2 cytokines were only produced by innate lymphoid cells and T cells **(Figure 2E and 2F)**. Monocytes showed spontaneous release of CCL2 and IL1β cytokines even in absence of stimulation, as described before^24^ **(Supplementary Figure 4)**.

**Figure 1.**
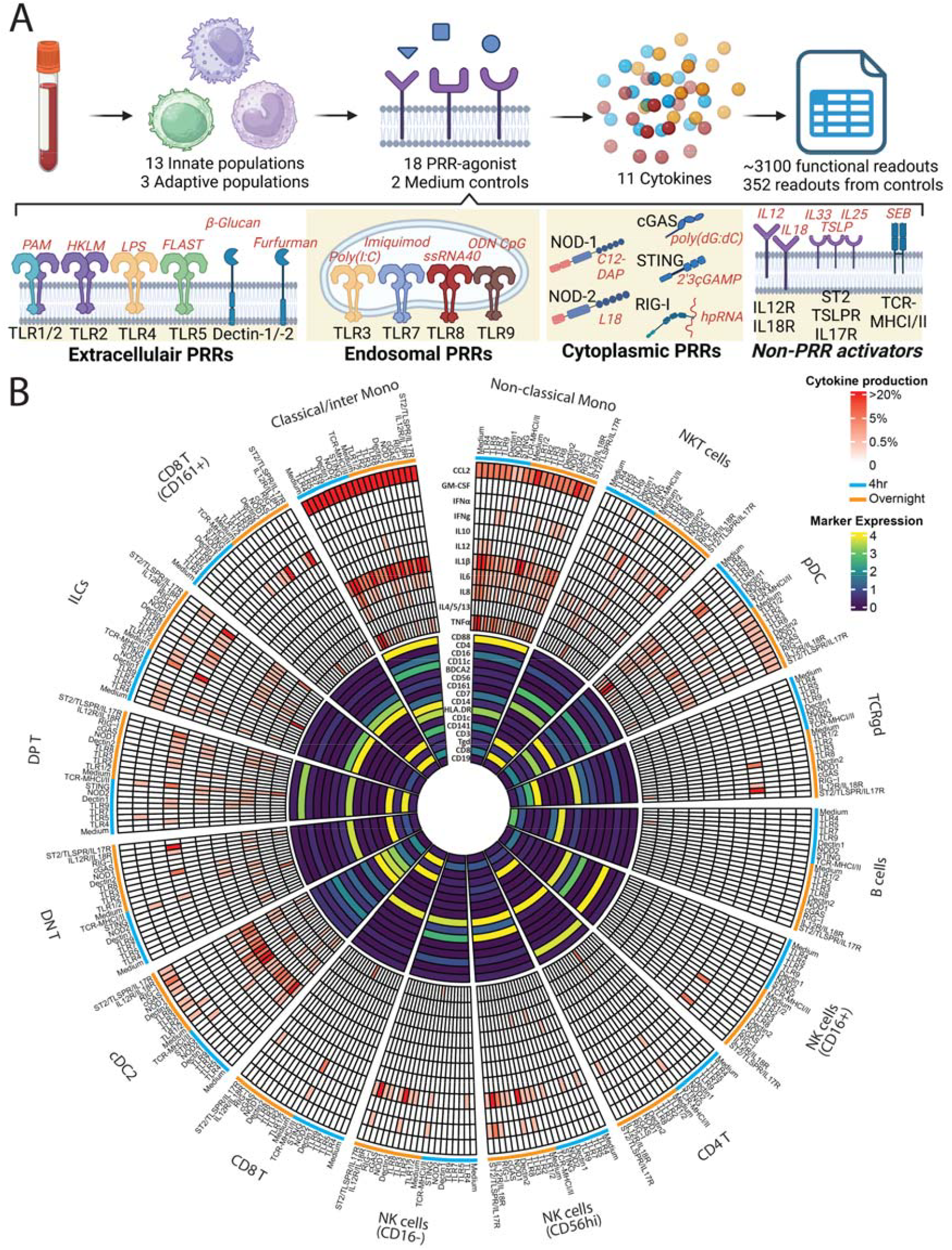
Quantification of innate and adaptive immune cell populations and their function against different pattern recognition receptor agonists. To study the response of innate populations to stimulation with different PRR-agonists, 18 peripheral mononuclear cell (PBMC) samples (cohort 1) were used to generate a holistic overview of the various PRR signalling pathways across the immune system. A) Overview of the functional assay whereby PBMCs were stimulated with 10 different PRR-agonists overnight and 8 different PRR-agonists for 4 hours. Production of 11 different cytokines by 16 different immune cell populations was measured for each stimuli, thereby generating ∼3100 readouts. B) Heatmap of marker expression is shown in the inner-circle for 16 different immune cell populations that were manually gated. Median intensity per population is shown after arcsinh transformation. For each population, the percentage of cytokine producing cells per cytokine in response to PRR agonists stimulated for 4 hours (blue) or overnight (orange), is shown in the outer-circle. The average of cytokine producing cells is shown for each population from cohort 1. In total, 462 combinations had an average cytokine production of >0.5%. *Abbreviations; PRR, pattern recognition receptor; TLR, Toll-Like Receptor; ILCs, Innate lymphoid cells; DP T, double-positive T cells (CD4+CD8+); DN T, double-negative T cells (CD4-CD8-); mono, monocytes; pDC, plasmacytoid dendritic cells; cDC, classical dendritic cells; NK cells, natural killer cells; NKT cells, natural killer T cells/*

**Figure 2.**
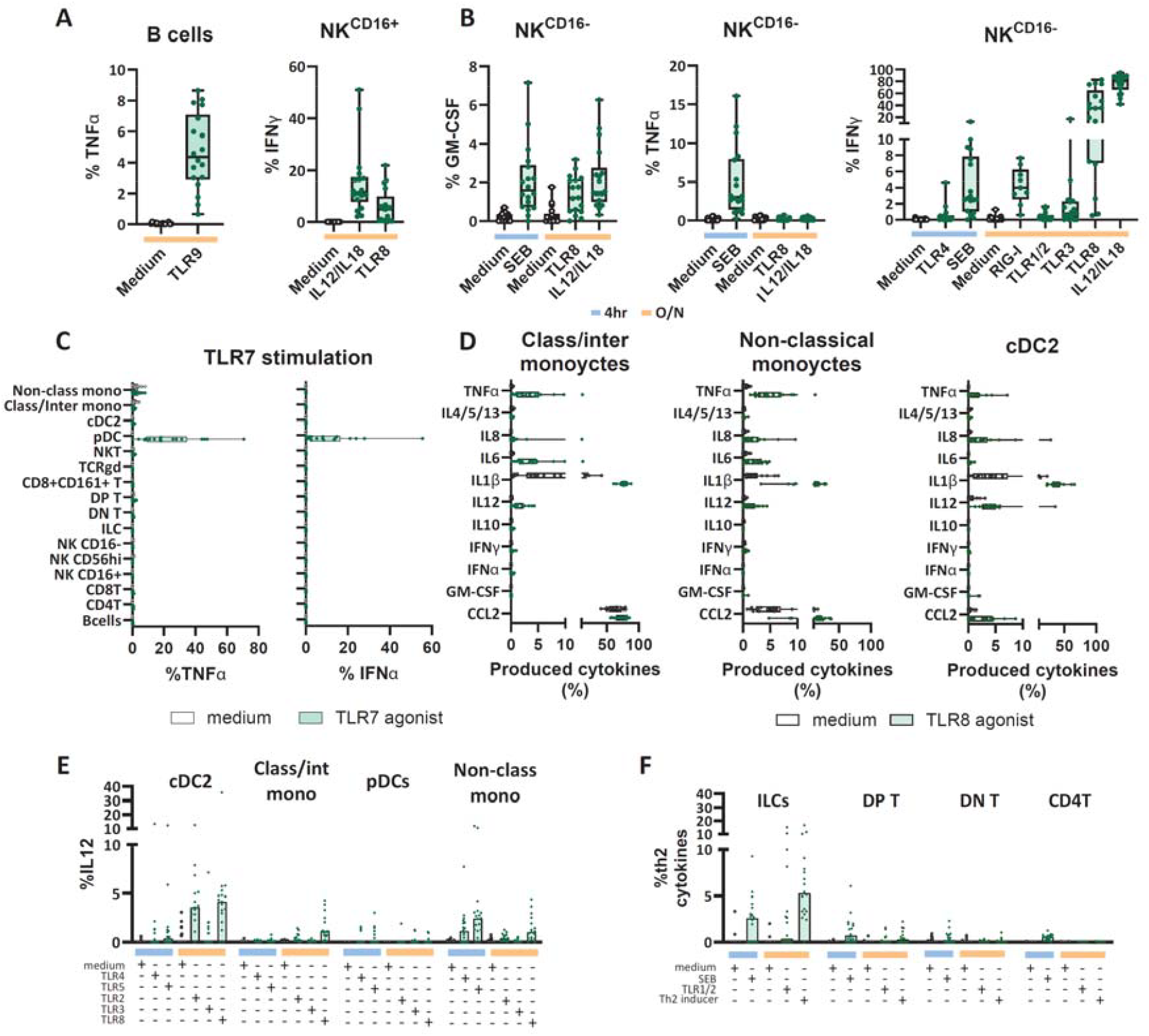
Innate and adaptive immune cell populations and their function against different pattern recognition receptor agonists. To study the response of innate populations to stimulation with different PRR-agonists, 18 peripheral mononuclear cell (PBMC) samples (cohort 1) were used to generate a holistic overview of the various PRR signalling pathways across the immune system. A and B) Shown are boxplots of populations that only respond to TLR8 and TLR9 stimulation (A) and populations that respond to multiple PRR agonists and produce different cytokines (B). Stimuli are coloured below the x-axis according to stimulation duration (blue; 4 hours, orange; overnight). C and D) Shown are boxplots of populations that solely respond to TLR7 stimuli (C) and different subsets of monocytes that produce multiple different cytokines in response to TLR8 stimulation (D). E and F). Boxplots showing which populations are responsible for IL12 production (E) or production of th2-cytokines (F) in response to different PRR agonists. *Abbreviations; TLR, Toll-Like Receptor; ILCs, Innate lymphoid cells; DP T, double-positive T cells (CD4*^*+*^ *CD8*^*+*^ *); DN T, double-negative T cells (CD4* ^*−*^ *CD8*^*−*^ *); mono, monocytes; pDC, plasmacytoid dendritic cells; cDC, classical dendritic cells; NK cells, natural killer cells*.

In summary, we were able to obtain, in a single assay, a holistic overview of the various PRR signalling pathways across the immune system, providing results congruent with data from literature^19-23, 25, 26^.

### Functionality of immune cell populations differ between people from different geographical areas

To compare the functional capacity of the immune system in people from rural and urban environments, we analysed PBMCs from rural Indonesians, urban Indonesians and healthy Europeans (n=6/group, **table 1**). Due to limited number of PBMCs available per sample, we prioritized 16/18 stimuli and compared the 420 functional readouts that had a frequency of >0.5% cytokine secreting cells across groups **(Supplementary Figure 3)**. Of these, 25 conditions were significantly different between the groups, with the rural Indonesians showing the most distinct profile based on hierarchical clustering **(Figure 3A)**. Of these 25 conditions, 6 were shared between the groups **(Figure 3B)**. In total, each of the 13 innate populations responded differently between groups against at least 1 PRR-agonist, with NK cells and monocytes showing most functional differences **(Figure 3A, Supplementary Figure 5A)**.

**Figure 3.**
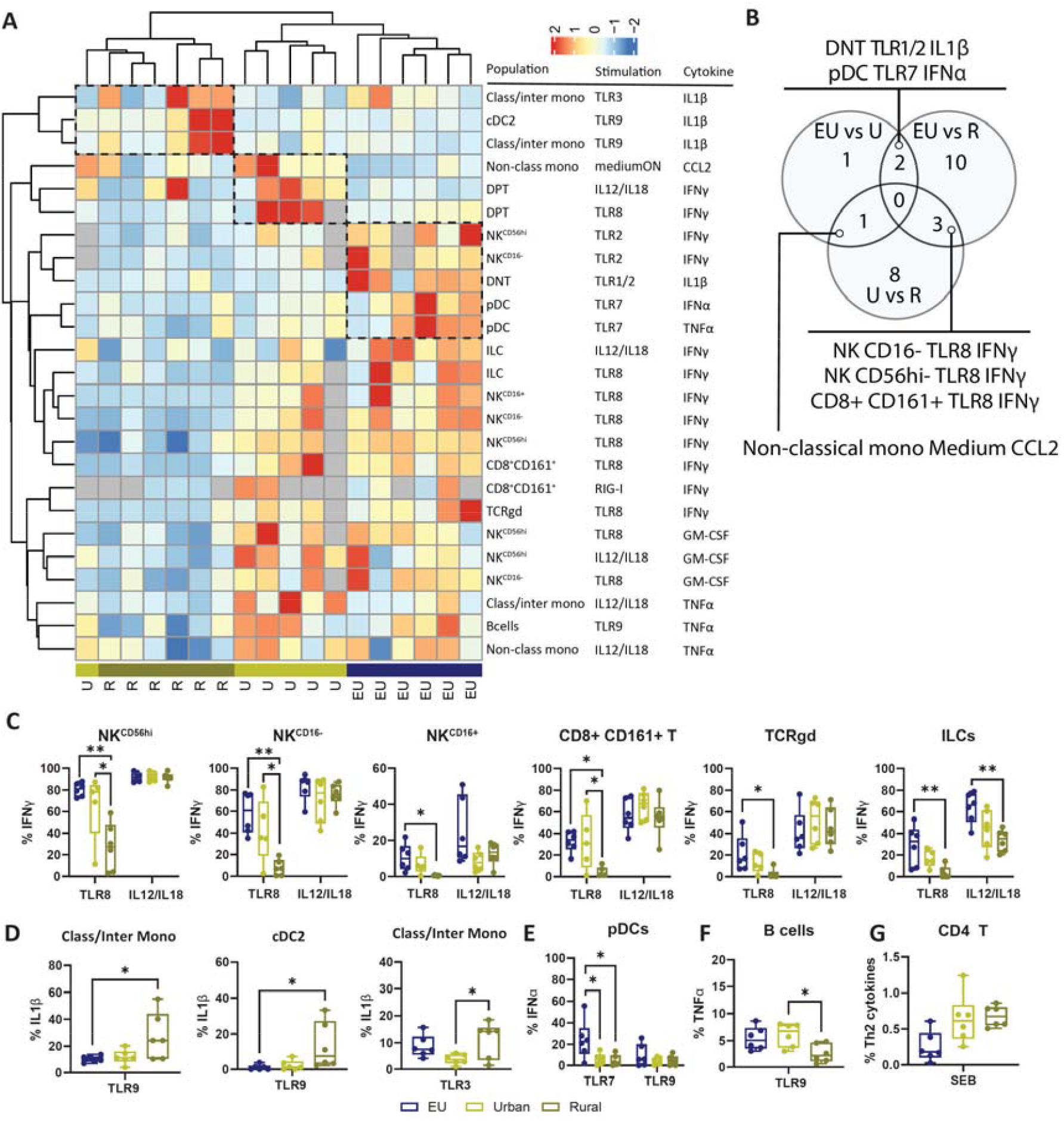
Functional differences of immune cell populations between people from different geographical areas. The function of the innate immune repertoire was investigated for people from different geographical areas by using the “function-omics” assay on PBMCs from rural Indonesians (R; gold), urban Indonesians (U; yellow) and healthy Europeans (EU;blue). A) Hierarchical clustered heatmap (Complete-linked, Euclidian distance) of z-scaled percentages of cytokine production for each of the 25 significant combinations (population/stimulation and produced cytokine; rows) of samples from each group (column). Grey-filled boxes indicate not performed. Dotted-line represents unique clusters per group. B) Venn-diagrams showing distribution and overlap of significant combinations between different groups. C,D,E,F and G) Shown are boxplots of immune populations that were functionally different between groups in response to different PRR agonists. *Abbreviations; PRR, pattern recognition receptor; TLR, Toll-Like Receptor; ILCs, Innate lymphoid cells; DP T, double-positive T cells (CD4*^*+*^ *CD8*^*+*^ *); DN T, double-negative T cells (CD4*^*−*^ *CD8*^*−*^ *); mono, monocytes; pDC, plasmacytoid dendritic cells; cDC, classical dendritic cells; NK cells, natural killer cells; NKT cells, natural killer T cells*

Striking differences in function were observed between innate lymphoid populations, such as NK cell subsets, CD8^+^CD161^+^ T cells, TCRgd cells and ILCs, which produced less IFNγ in response to TLR8 stimulation in rural Indonesians compared to urban Indonesians and Europeans (Figure 3C). The difference in IFNγ production was not due to intrinsic impairment of the innate lymphocytes, since no difference in response to IL12/IL18 was found for all these populations with the exception of ILCs. Moreover, no statistically significant differences were observed in response to other endosomal PRR agonists, although a trend for a lower response to TLR3 was observed for rural Indonesians **(Supplementary Figure 6)**. Contrary to the decreased responses of the lymphoid compartment in rural Indonesians, a more pro-inflammatory response from cDC2s and classical/intermediate monocytes was observed, with significantly more IL1β production in response to TLR3 and TLR9 stimulation **(Figure 3D; Supplementary Figure 5B)**. cDC2s also displayed a trend for increased IL1β production upon TLR8 stimulation, which was negatively correlated with the frequency of IFNγ producing lymphoid cells **(Supplementary Figure 7)**. There were also differences between urban Indonesians and Europeans, for example pDCs from both rural and urban Indonesians produced significantly lower frequencies of IFNα and TNFα in response to TLR7 than Europeans **(Figure 3E)**. No difference in response to TLR9 was observed from pDCs. Of the adaptive immune cell populations, B cells produced less TNFα upon TLR9 sensing in the rural Indonesian group compared to the other groups **(Figure 3F)**. Furthermore, a non-significant trend was observed for CD4 T cells to produce more Th2-cytokines in the rural Indonesian group in response to SEB stimulation, in line with literature^12^ (Figure 3G).

An independent cohort (Table 1; cohort 2) was used to investigate whether altered TLR levels could underlie the distinct functional responses between groups. No significant differences were observed for TLR levels among innate lymphoid-derived cells, while increased TLR7, TLR8 and TLR9 expression were observed for myeloid populations of rural Indonesians, suggesting that alterations in responses were not determined by receptor abundances but likely due to alterations in downstream pathways **(Supplementary Figure 8)**.

In conclusion, a more pro-inflammatory state of myeloid immune populations, alongside increased expression of endosomal TLR sensors was observed in rural Indonesians, while innate lymphoid populations showed decreased interferon production after TLR8 stimulation.

### Innate baseline response to TLR8 stimulation predicts immunogenicity of SARS-CoV-2 mRNA vaccination

We next wanted to investigate if the functional capacity of immune cells to endosomal TLRs (TLR3, 7, 8 and 9) and cytosolic sensors (cGAS, STING and RIG-I) could predict primary SARS-CoV-2 mRNA-1273 vaccination responses, as the underlying sensing mechanisms that engage the innate immune system for this relatively novel vaccine type are still partly unknown in humans^27^. We measured the baseline functional capacity of PBMCs of 18 SARS-CoV-2 infection- and vaccine-naïve European individuals (Table 1; cohort 3) and associated this with spike-specific antibody titers and spike-specific T and B cells^17^ 2 weeks (day 43) after the second dose of vaccination **(Figure 4A, Supplementary Figure 9A)**. Spike-specific CD4^+^ T cells, B cells and antibodies were all correlated with each other **(Supplementary Figure 9B)**, whereas spike-specific CD8^+^ T cells showed a distinct profile and did not correlate with the antibody production. When looking at baseline frequencies of immune cell populations, and not their functional capacity, no significant correlations (p<0.01 and r>0.5) of immune population frequencies could be found with Spike-specific antibody titers or spike-specific T and B cells **(Figure 4B)**. This highlights the complexity of linking cell frequencies to vaccination responses. However, when assessing functional capacity, significant positive correlations (p<0.01 and r>0.5) were found for different TLR8 responding NK subsets and RIG-I responding ILCs, with the percentages of spike-specific B cells and spike-specific CD4^+^ T cells after vaccination, respectively **(Figure 4C and 4D)**. In contrast, different subsets of IL1β producing monocytes and cDC2s in response to agonists for TLR2, TLR8 and RIG-I negatively correlated with the IgG^+^ spike-specific serum antibody titers detected in plasma after vaccination **(Figure 4C and 4D)**.

**Figure 4.**
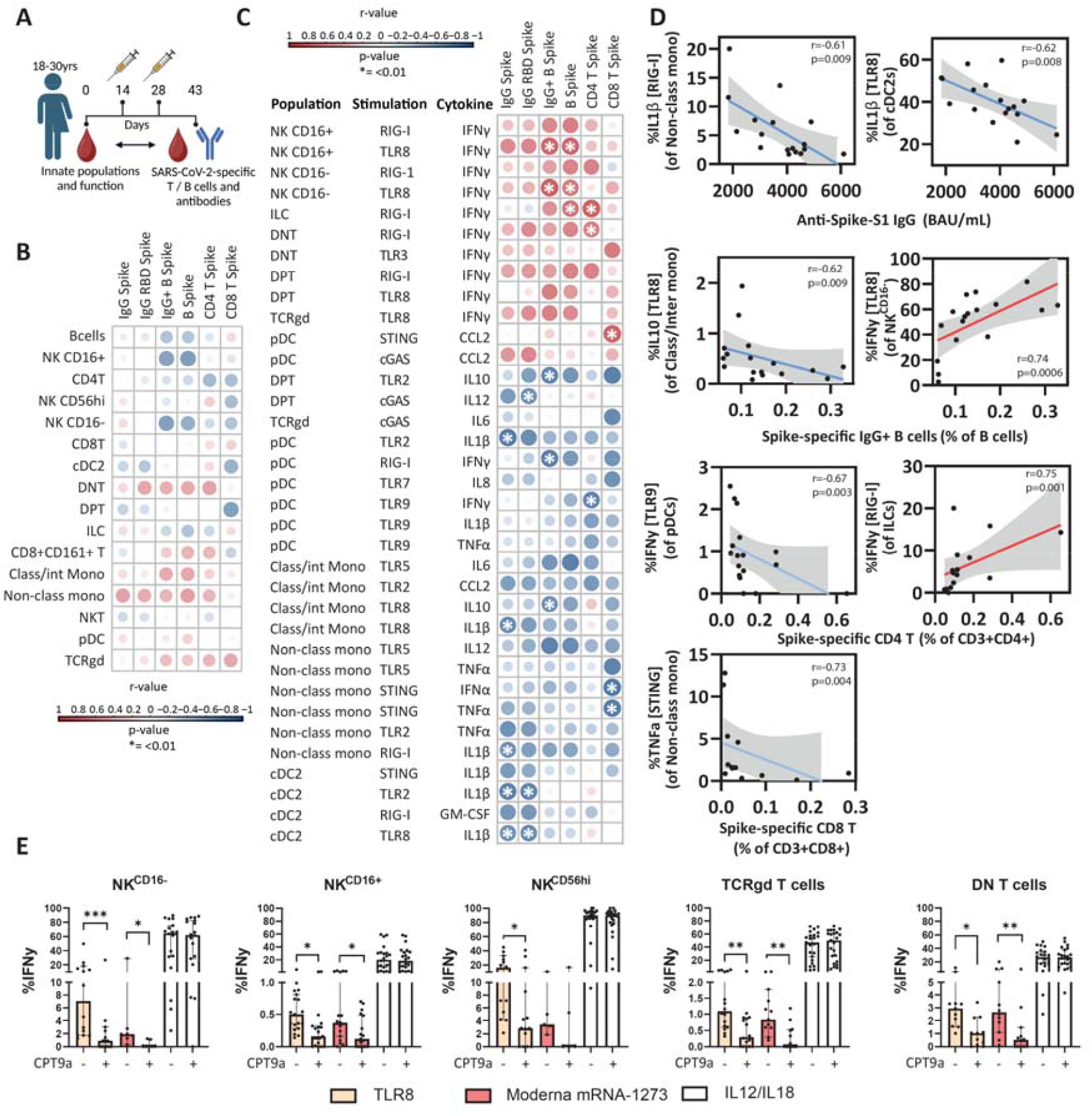
Innate baseline functional response to PRR stimulation predicts immunogenicity of SARS-CoV-2 mRNA vaccination. The baseline functional capacity of PBMCs of 18 SARS-CoV-2 naïve European individuals was measured with priority on reactivity against endosomal TLR agonists (TLR3,7,8 and 9) and cytosolic sensors cGAS, STING and RIG-1. The functional response was then compared to the frequencies of SARS-CoV-2-specific T and B cells and titers of SARS-CoV-2-specific antibodies. A) Overview of the study where 2 doses of moderna mRNA-1273 vaccination were administered, and peripheral blood was obtained before vaccination and 2 weeks after second dose. B and C) Correlation heatmap reporting Spearman correlation coefficients (r) and P values for each comparison. Cut-off for significant correlations are r>0.5 and p<0.01 and are shown as white asterix. Baseline frequencies (B) or functional responses (C) of immune cell populations are compared to frequencies of SARS-CoV-2-specific T or B cells or titers of SARS-CoV-2-specific antibodies. D) Representative Spearman correlation plots are shown for immune cell populations where the function correlated with the vaccine immunogenicity. E) PBMCs were stimulated with TLR8 agonist ssRNA40, Moderna mRNA1273 and IL12/IL18, with and without TLR8 inhibitor CPT9a. Statistical differences were assessed with Wilcoxon signed-rank tests. \**P* < 0.05; \*\**P* < 0.01; \*\*\**P* < 0.001. Bar plots depict median with 95%CI *Abbreviations; PRR, pattern recognition receptor; TLR, Toll-Like Receptor; ILCs, Innate lymphoid cells; DP T, double-positive T cells (CD4*^*+*^ *CD8*^*+*^ *); DN T, double-negative T cells (CD4*^*−*^ *CD8*^*−*^ *); mono, monocytes; pDC, plasmacytoid dendritic cells; cDC, classical dendritic cells; NK cells, natural killer cells; NKT cells, natural killer T cells*

The association between TLR8 responses and mRNA-1273 immunogenicity might indicate direct sensing of mRNA through this pathway, which was surprising given the replacement of uridine nucleotides with naturally occurring derivatives of uridine that serve to reduce innate sensing within mRNA vaccines^28^. To assess whether residual TLR8 sensing of mRNA vaccines still occurs, we stimulated PBMCs *in vitro* with mRNA-1273 with and without TLR8 blocking (CPT9a). Stimulation with mRNA-1273 led to production of IFNγ by the same innate lymphoid populations that we previously found to be predictive for a good vaccine response (CD16^+^ NK and CD16^-^ NK cells), which could be blocked by inhibiting TLR8 sensing with CPT9a **(Figure 4E)**. As controls we confirmed that TLR8-induced, but not IL12/18-induced, IFNγ production was reduced by TLR8 blocking. Classical/intermediate and non-classical monocytes produced IL1β in response to mRNA-1273, but this was not affected by TLR8 blockade **(Supplementary Figure 10)**. This indicates a different sensing pathway was responsible for the inflammasome activation following mRNA-1273, likely in response to the lipid nanoparticle^28^.

In summary, high innate lymphoid-derived IFNγ responses to intracellular nucleic acid sensor stimulation, including TLR8, associated with good CD4^+^ T cell, B cell and antibody responses to mRNA-1273 vaccination.

### *in vitro* stimulation with mRNA vaccines induces reduced IFNγ production in rural Indonesians

As we observed that TLR8 is critical for responses to mRNA vaccines, and this pathway was reduced in rural Indonesians, we hypothesized that PBMC from rural Indonesians would show reduced responses to mRNA stimulation. Therefore, we stimulated a fourth independent cohort of PBMCs of Europeans and rural Indonesians *in vitro* with Comirnaty mRNA-XBB (mRNA-1273 was no longer available at this point; Table 1; cohort 4). First, we confirmed that also in this second cohort TLR8-induced innate lymphocyte responses were reduced in rural Indonesians **(Figure 5A)**. Moreover, IFNγ production by CD16^-^ NK innate lymphocytes upon mRNA-XBB stimulation was found to be lower in rural Indonesians compared to Europeans. Classical/intermediate monocytes also displayed a trend for increased IL1β production upon TLR8 stimulation, which, together with cDC2s, negatively correlated with the frequency of IFNγ producing CD16^-^ NK cells **(Figure 5B)**. IFNγ production in response to Comirnaty mRNA-XBB by CD16^-^ NK cells and TCRgd T cells, correlated positively with the TLR8 response **(Figure 5B)**.

**Figure 5.**
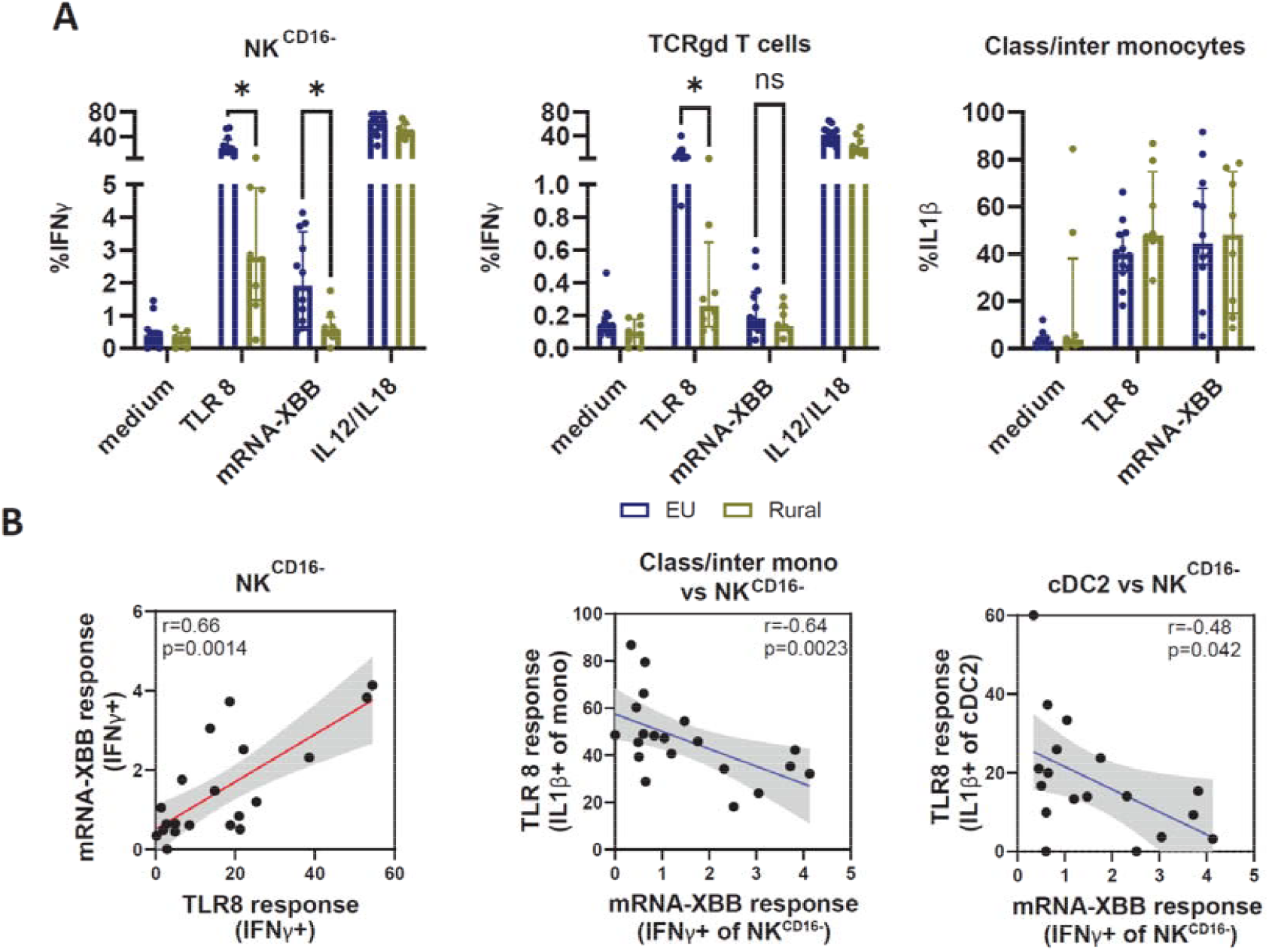
Response to SARS-CoV-2 Comirnaty mRNA-XBB vaccine *in vitro* of rural Indonesians and Europeans. The response to mRNA vaccines of the innate immune repertoire from rural Indonesians and Europeans was investigated by *in vitro* stimulation with Comirnaty mRNA-XBB and was compared to the response to TLR8. **A)** Shown are boxplots of immune populations that were functionally different between groups in response to Comirnaty mRNA-XBB and TLR8. **B)** Spearman correlation plots are shown for immune cell populations where the response to TLR8 correlated with the Comirnaty mRNA-XBB vaccine immunogenicity. Statistical differences were assessed with multiple Mann-Whitney U tests, FDR corrected. \**P* < 0.05. Bar plots depict median with 95%CI. Spearman correlation coefficients (r) and P values for each comparison are depicted in the correlation plots.

Thus, we confirmed reduced TLR8-induced innate lymphocyte responses in an independent cohort of rural Indonesians, which correlated with reduced responses upon *in vitro* stimulation with a mRNA vaccine. Moreover, a strong inflammasome response was linked with impaired responses of innate lymphoid populations to mRNA vaccines.

## Discussion

In this study, we developed a holistic approach to study the functional capacity of the innate immune system to respond to stimulation of different pattern recognition receptors. We showed that a more pro-inflammatory state of myeloid immune cells is observed in rural Indonesians. The inflammasome response of cDC2 and monocytes to TLR8 correlated negatively with the decreased function of innate lymphoid-derived cells towards mRNA vaccines, as well as decreased immunogenicity. Furthermore, we showed that interferon responses by innate lymphoid populations positively correlates with IgG+ spike-specific antibody titers and spike-specific CD4+ T cells after vaccination.

We showed that *in vitro* stimulation with mRNA-1273 led to IFNγ production by innate lymphoid populations in a TLR8-dependent manner. TLR8 was previously not identified as an important pathway in a study where mice were vaccinated with BNT162b2, as mice do not have functional TLR8^29^. The modification of the nucleic acids in the mRNA vaccine should prevent sensing by TLR8^30, 31^, but our data indicate that this reduction is not complete in human-derived immune cells.

Predictions for a good vaccine response often only rely on the produced antibodies. Here we not only looked at the produced antibodies but also at the whole cellular response and memory formation of B and T cells. While antibodies, B cells and CD4^+^ T cells responses showed good correlations with each other and the functional pathways that were predictive of them, CD8^+^ T cells were not associated with them. In line with the separate mode of action of CD8^+^ T cell induction, mice infected with hookworms demonstrated compromised spike-specific T cell responses, although this did not affect antibody responses^32^. Thus different response pathways are associated with CD8^+^ T cell immunity compared to humoral immunity, and this increased granular understanding has implications for 1) vaccines against pathogens for which a good CD8+ T cell response is wanted, or 2) for individuals that are not able to generate adequate antibodies.

The reasons behind vaccine hyporesponsiveness for individuals from LMIC are still poorly understood since various factors may contribute to this reduced immunogenicity^10^. In general, few studies highlight the role of an increased pro-inflammatory transcriptomic profile, which was associated with reduced vaccine responses^5, 6, 8^. However, these studies were not able to pinpoint which cells displayed this pro-inflammatory profile and if that would impact specific PRR pathways, nor did they investigate this in a LMIC setting. Our approach allowed us to investigate this further, where in rural Indonesians we found a strong pro-inflammatory cDC2 response towards TLR8 and TLR9, which could specifically impact the Natural Killer-Dendritic Cell immune axis, which is important in the initiation and coordination of adaptive immunity^33, 34^. Indeed, IL1β production by cDC2s was negatively associated with IFNγ production by innate lymphoid populations upon TLR8 and Comirnaty mRNA-XBB stimulation. Whether this can be fully attributed to the inflammasome itself remains difficult, since it has also been reported that IFNγ is a crucial regulator of inflammasome activation^35, 36^.

Higher IFNγ production upon TLR8 stimulation was associated with improved *in vivo* responses to SARS-CoV-2 in Europeans. Together with the reduced IFNγ responses upon TLR8 or mRNA vaccine stimulation in rural Indonesians, this raises the hypothesis that mRNA vaccines might suffer from vaccine hyporesponsiveness when deployed in LMIC. While there is no comparative *in vivo* data yet of mRNA vaccination in individuals from LMIC compared to high income countries, a clinical trial with the mRNA-1644 HIV vaccine is underway in Rwanda and South-Africa (NCT05414786). Another vaccine for which TLR8 was found to be important, the Rotavirus vaccine Rotarix^37^, has been shown to be one of the vaccines that also suffers from decreased efficacy in LMIC.

There are limitations to our study. It is known that the duration of *in vitro* stimulation impacts the cytokines that are released. Therefore, a limitation of our approach is that we are only measuring a snapshot (4 and 20hrs) of cytokine production. These different stimulation durations for the PRR agonists make it furthermore difficult to compare them to each other. Also, in our approach we specifically stimulate only single PRR pathways, while it is known that stimulation of multiple different pathways simultaneously^38^ or in combination with Fcγ receptors^39^ can lead to synergistic responses. However, our parsimonious approach allows us to identify specific pathways that may be crucial in responses. Finally, stimulating each sample with 20 different conditions requires a large enough number of cells that may not be available in all settings, for example paediatric studies.

In summary, our study reveals key insights into the interplay between innate immune system responses, geographical variation, and vaccination outcomes, highlighting the predictive potential of baseline immune parameters and the critical role of specific PRR pathways in optimising vaccine design and addressing global disparities in immunogenicity.

## Acknowledgements

This work was supported by Leiden University Funds (LUF) awarded to Simon Jochems. This study was additionally supported by grants from the European Research Council (ERC) via the ERC Advanced Grant ‘REVERSE’ awarded to Maria Yazdanbakhsh (Grant No: 101055179), The Royal Netherlands Academy of Arts and Science (KNAW), Ref 57-SPIN3-JRP and Universitas Indonesia (Research Grant BOPTN 2742/ H2.R12/HKP.05.00/2013.), crowdfunding (Wake Up to Corona; a crowdfunding for COVID-19-related research set up by Leiden University, the Netherlands), applications through Bontius stichting Leiden (BS140-20210186), U-Needle (developer and producer of the Bellamu® needle) and two Dutch philanthropic organizations (SAL stichting Apothekers and Diraphte). The sponsors are nonprofit organizations that support science in general. They had no role in gathering, analyzing, or interpreting the data. The authors gratefully acknowledge the flow cytometry core facility (FCF) at LUMC, Leiden, the Netherlands for technical support regarding spectral flow cytometry.

## Supplementary Figures and Tables

**Supplementary Figure 1.**
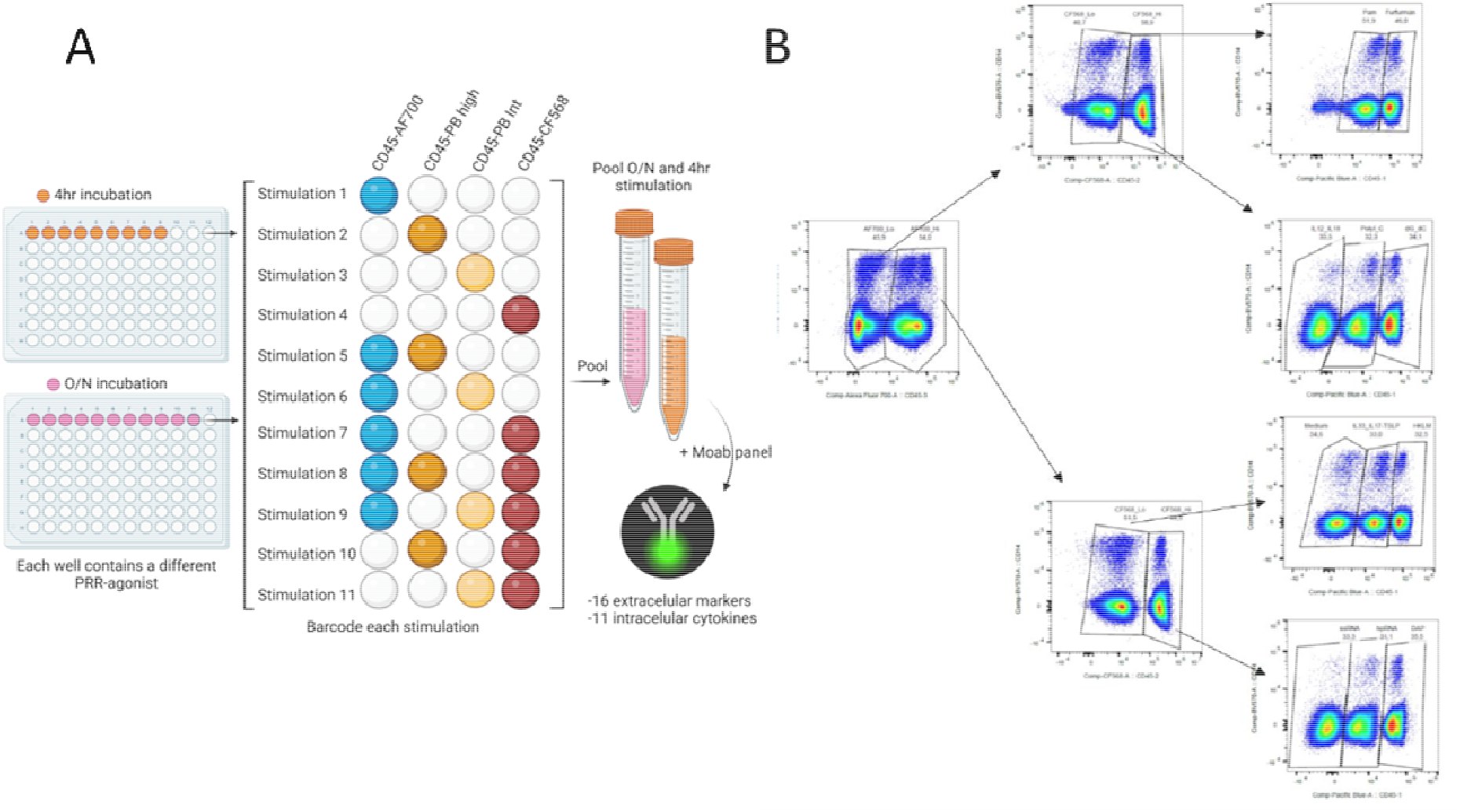
Barcoding of PRR-agonist stimulated immune cells. **A)** Peripheral blood mononuclear cells (PBMCs) from each donor were split over 9 wells for 4 hour stimulation (orange) and 11 wells for overnight stimulation (pink) to stimulate with 18 different PRR-agonists and 2 medium controls. After stimulation, each well from each plate (4 hours or overnight) was barcoded according to the schematic using CD45-AF700, CD45-PB^high^, CD45-PB^int^ and CD45-CF568. Barcoded wells were then pooled and stained using monoclonal antibodies for extracellular markers and after fixation and permeabilization with intracellular markers. B) After acquisition of the pooled and barcoded samples (4hours and overnight), each barcoded stimulation was manually gated and exported. Shown is a representative pooled sample containing overnight stimulated conditions. *Abbreviations; PRR, pattern recognition receptor; AF700, AlexaFluor700; PB, PacificBlue; CF568, Cyanine-based Fluorescent 568*

**Supplementary Figure 2.**
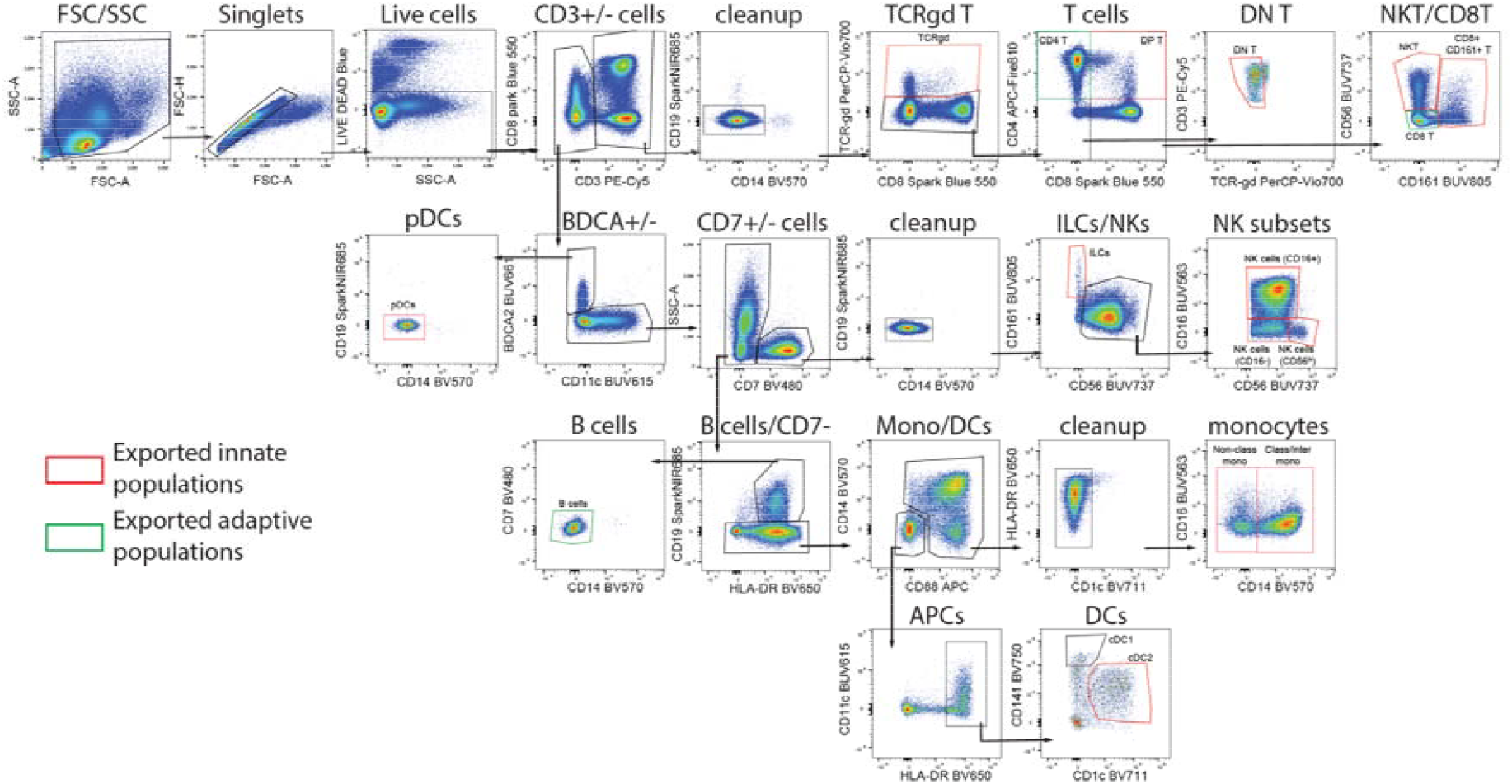
Manual gating and exporting of immune cell populations after stimulation. For gating innate and adaptive immune cell populations after stimulation, Peripheral Blood Mononuclear Cells (PBMCs) were thawed, stimulated with 18 PRR-agonists, barcoded using different CD45 monoclonal antibodies and stained with a cocktail of antibodies targeting extracellular proteins. Single-cell live lymphocytes were first gated before the CD3+ lymphocytes were separated. From the CD3-subset pDCs were gated and CD3-CD7+ lymphocytes were separated from the myeloid lineage. Innate (red gate) and adaptive (green gate) immune cell populations were exported for further analyses. Intermediate (CD14+CD16+) and classical (CD14+CD16-) monoyctes were exported as one mixed population due to downregulation of CD16 after stimulation. *Abbreviations; ILCs, Innate lymphoid cells; DP T, double-positive T cells (CD4*^*+*^ *CD8*^*+*^ *); DN T, double-negative T cells (CD4*^*-*^*CD8*^*-*^*); mono, monocytes; pDC, plasmacytoid dendritic cells; cDC, classical dendritic cells; NK cells, natural killer cells; NKT cells, natural killer T cells; APC, Antigen presenting cells; BDCA2, blood dendritic cell antigen 2*

**Supplementary Figure 3.**
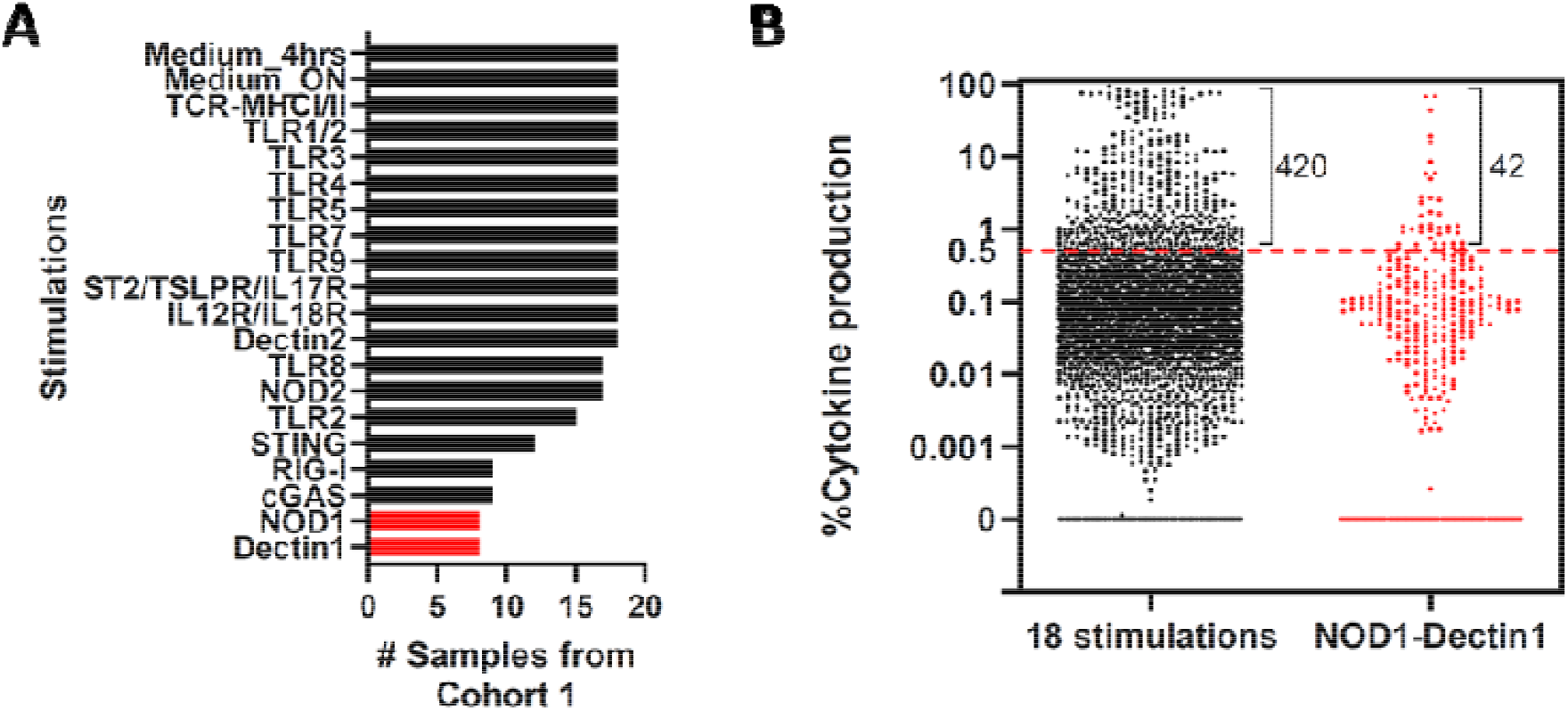
Number of samples that were stimulated per PRR agonist and number of cytokine producing conditions. **A)** Shown are the number of samples for each stimulation of cohort 1. For comparisons between groups a cut-off of at least 9 samples was selected. Stimulations in red indicate stimulation that did not fulfil this criteria. B) Dotplots whereby each dot represents the average cytokine production per cytokine (n=11) of each population (n=13) for every stimulation. In total 462 conditions showed more than 0.5% cytokine production. Stimulations for NOD1 and Dectin1 were dropped when differences between groups of cohort 1 were compared due to limited numbers of samples.

**Supplementary Figure 4.**
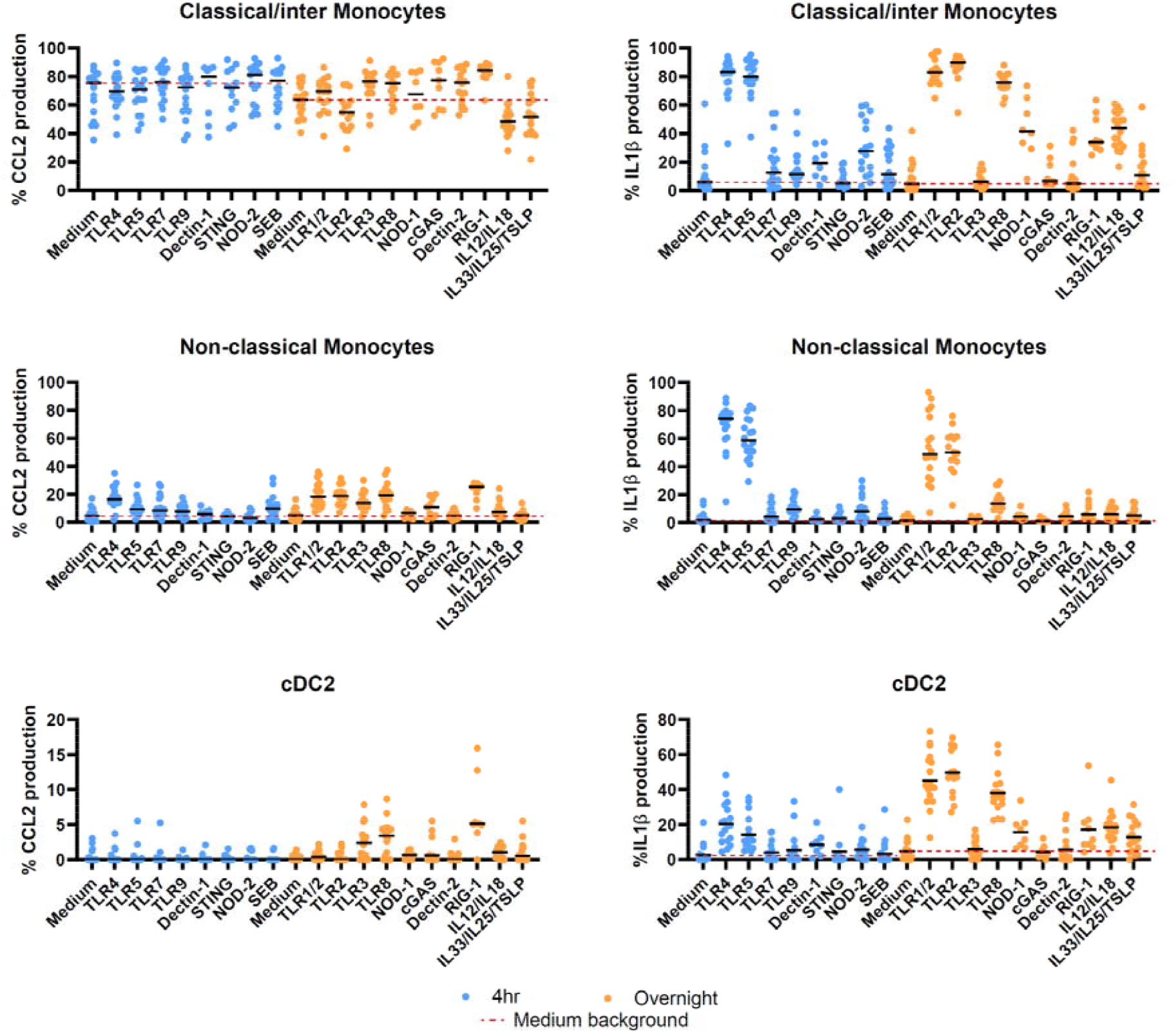
Spontaneous release of CCL2 and IL1β by monocytes and cDCs. The IL1β and CCL2 production of monocytes and cDC2s from 18 peripheral mononuclear cell (PBMC) samples from cohort 1 are depicted in response to different PRR-agonists. Stimuli are coloured according stimulation duration (blue; 4 hours, orange; overnight). Red dotted lines indicate medium background for 4 hour and overnight stimuli separately. *Abbreviations;TLR, Toll-Like Receptor; cDC, classical dendritic cells; NK cells*,

**Supplementary Figure 5.**
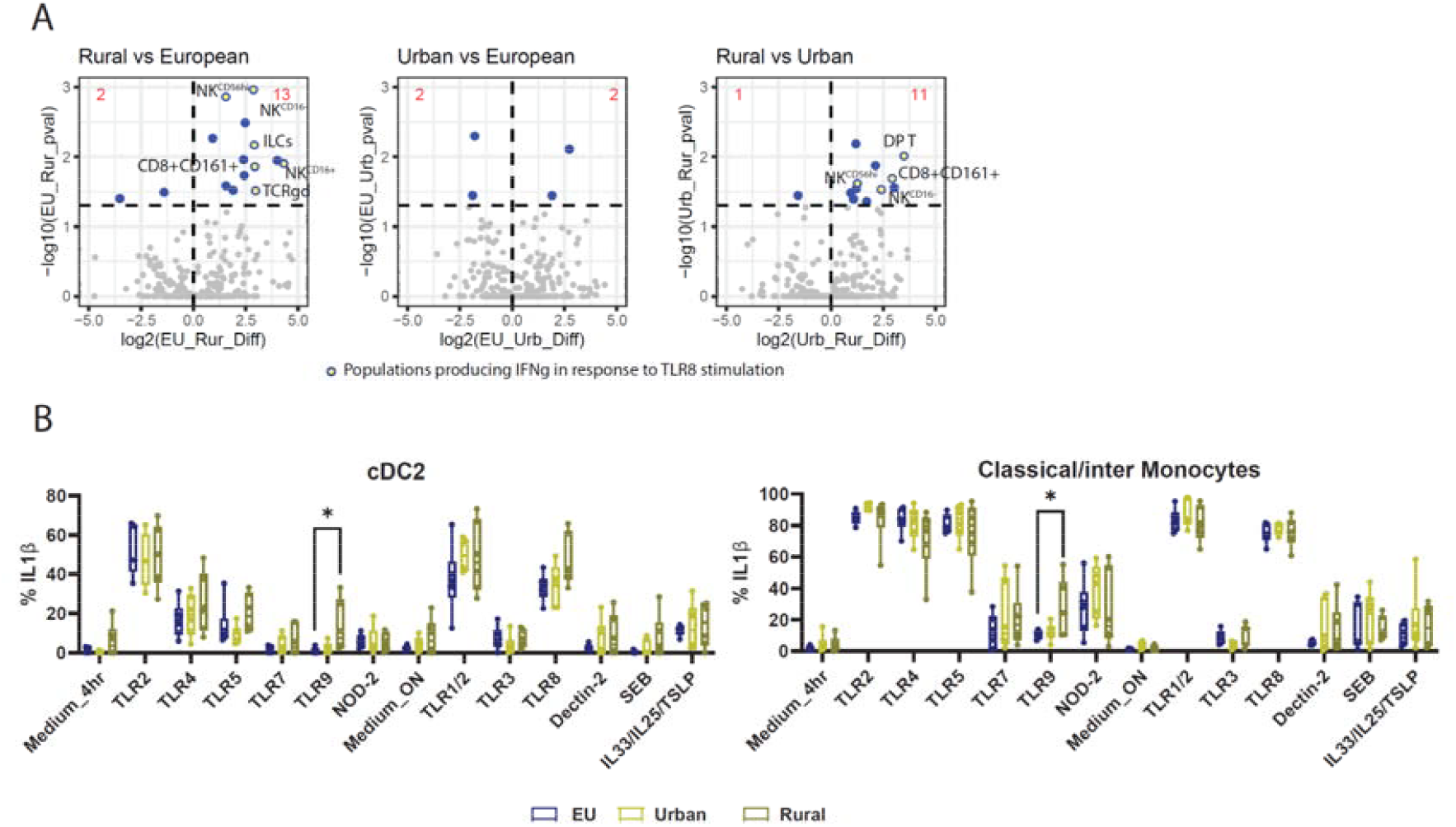
Functional differences of immune cell populations between people from different geographical areas. **A)** Volcano plots of 420 cytokine producing (>0.5%) immune cell populations in response to different PRR agonists between different study groups. Populations that produce IFNγ in response to TLR8 stimulation are highlighted by yellow filled circles. B) Boxplots of production of IL1β of cDC2s and classical/intermediate monoyctes in response to different PRR-agonists.

**Supplementary Figure 6.**
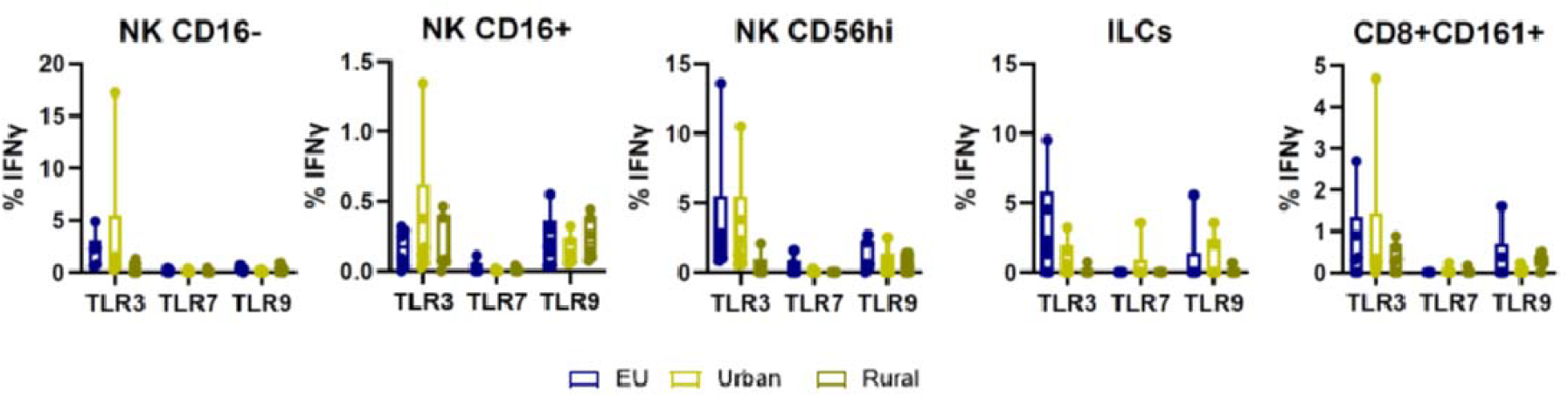
Functional response to endosomal TLR agonists by innate immune cell populations. Boxplots of production of IFNγ of innate lymphoid-derived immune cell populations (cohort 1) in response to different endosomal PRR-agonists.

**Supplementary Figure 7.**
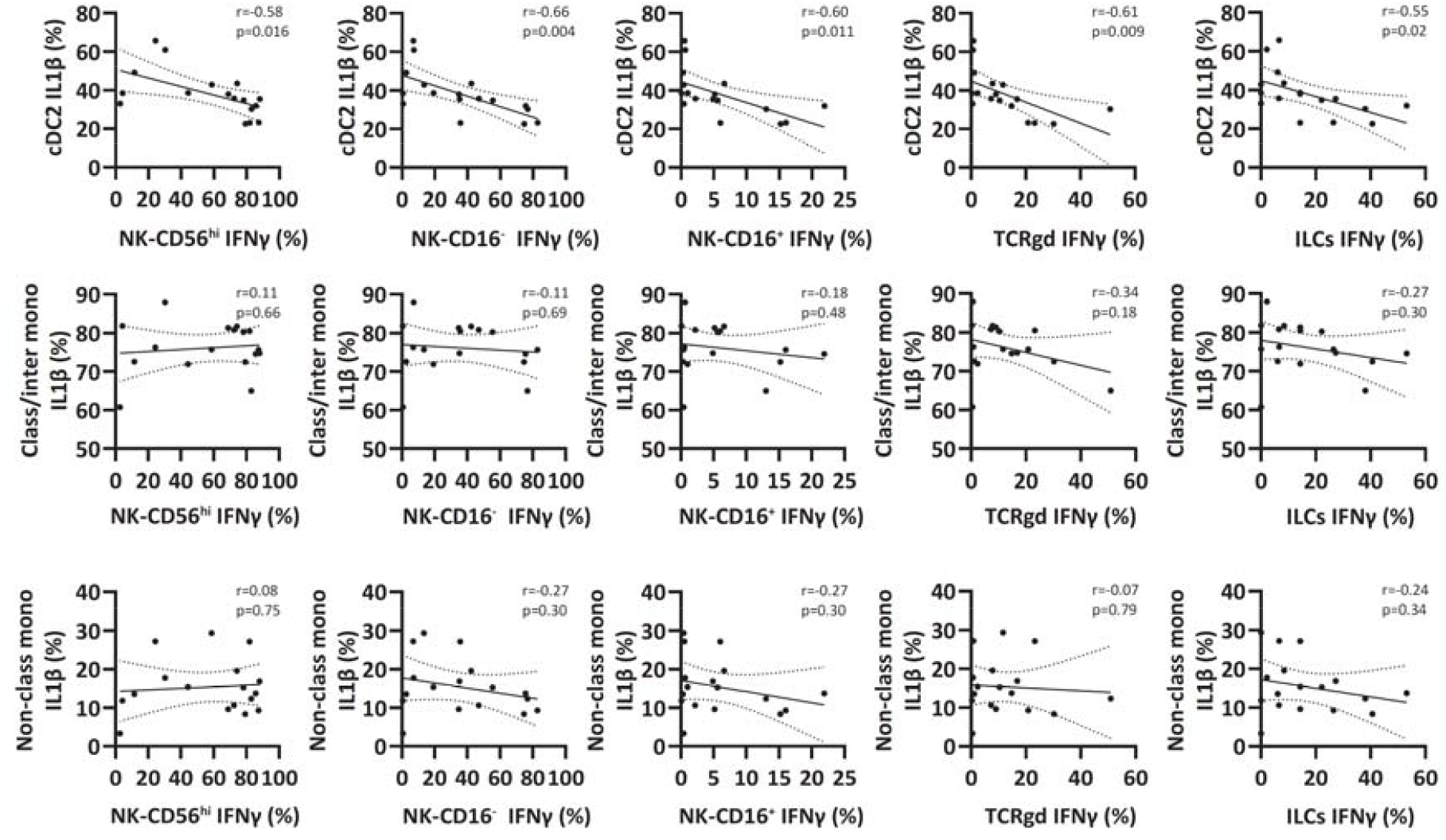
Correlation plots of TLR8-agonist responding myeloid and lymphoid-derived innate populations. Correlation plots are shown for IFNγ producing lymphoid-derived innate populations against IL1β producing cDC2s, (top row) IL1β producing class/intermediate monocytes (middle row) and IL1β producing non-class monocytes (bottom row). Spearman correlation coefficients (r) and P values for each comparison are shown within each plot.

**Supplementary Figure 8.**
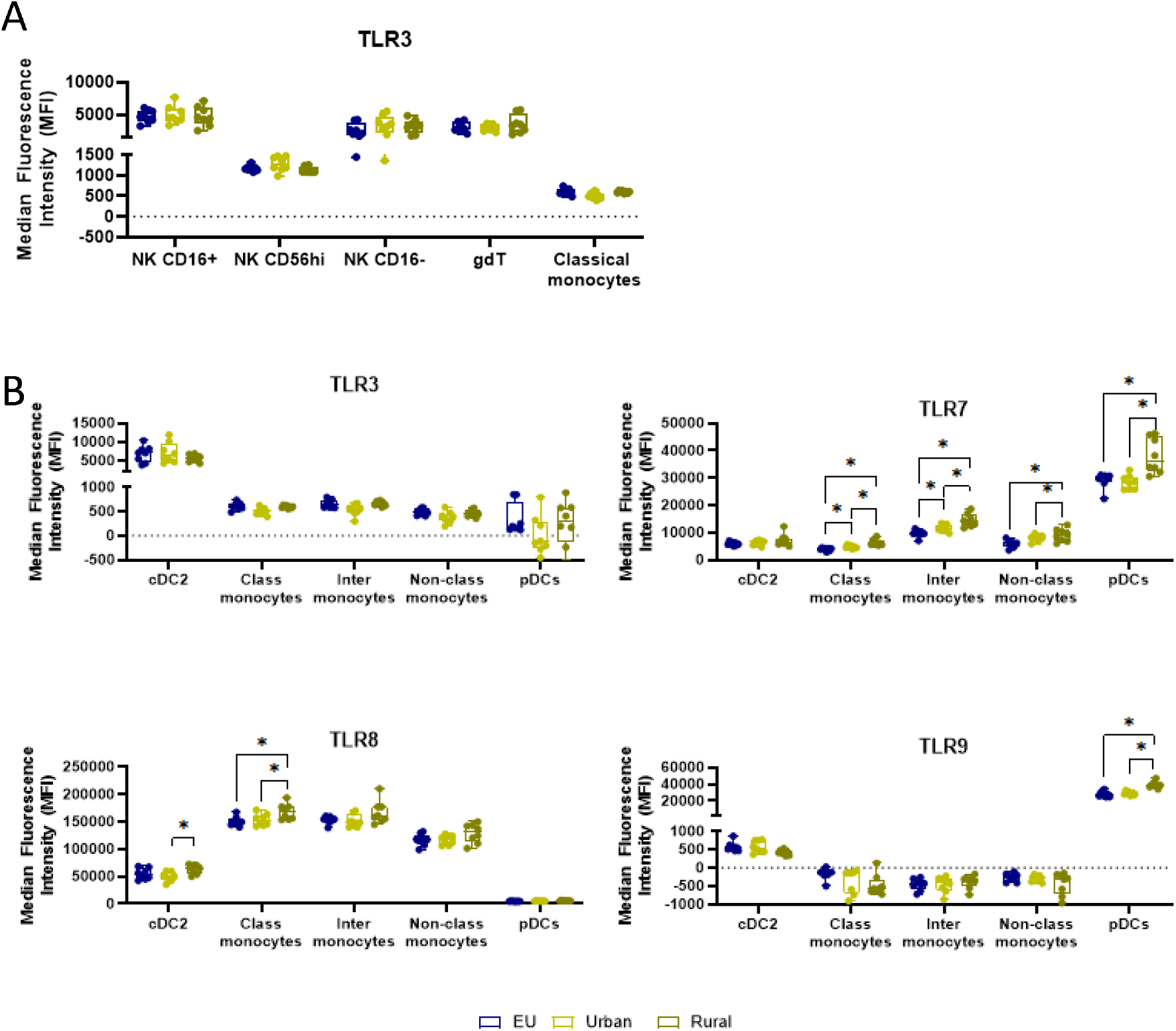
TLR-expression of innate immune cell populations. PBMCs from cohort 3 were used to measure the TLR expression of different lymphoid-derived innate immune cell populations and myeloid-derived innate immune cell populations. A) Expression of TLR3 is shown for different lymphoid-derived innate immune cell populations. B) Expression of endosomal TLRs (TLR3, TLR7, TLR8 and TLR9) is shown for different myeloid-derived innate immune cell population. Statistical differences were assessed with One-way Anova with FDR correction for multiple testing. \**P* < 0.05; \*\**P* < 0.01; \*\*\**P* < 0.001. Floating box plots depict median with min-max

**Supplementary Figure 9.**
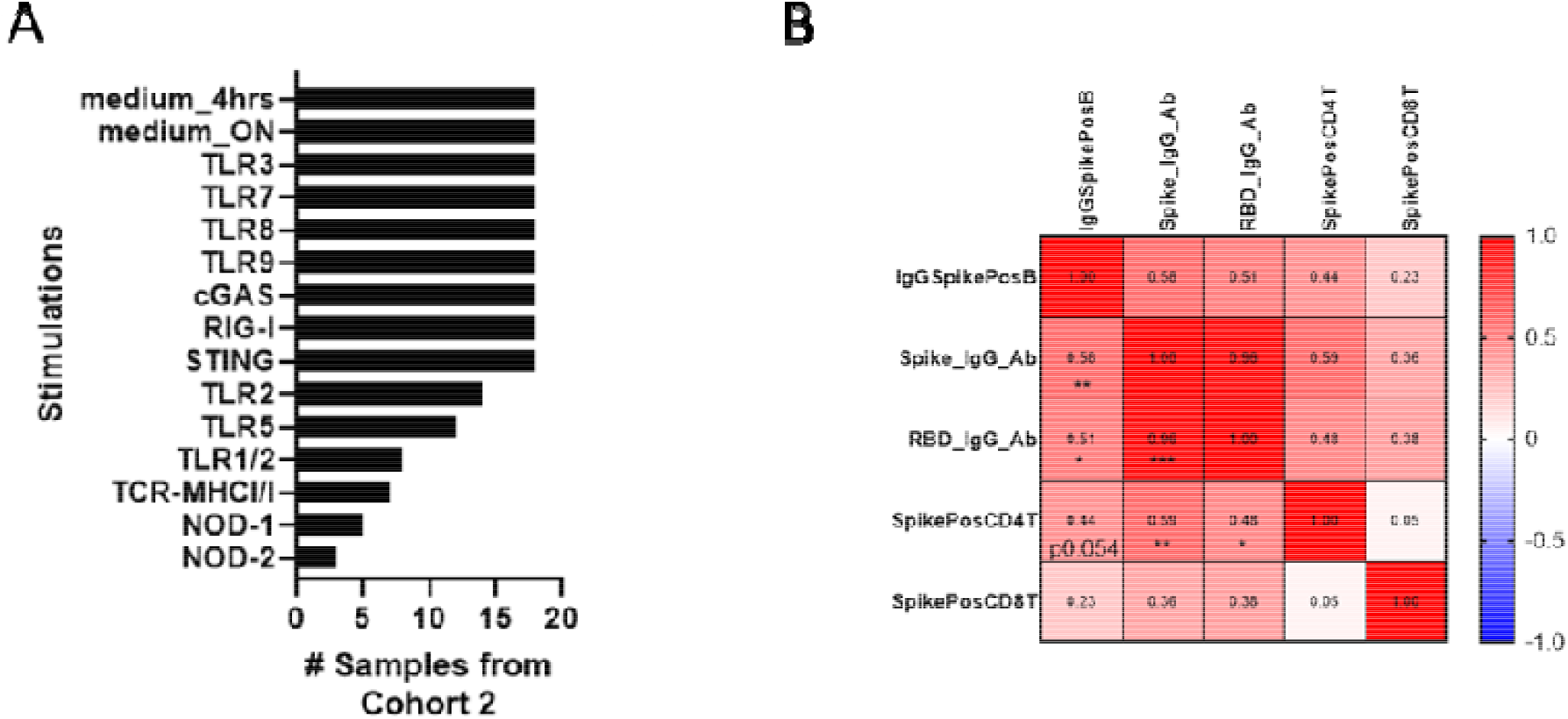
Overview of stimulations from cohort 2 and correlation matrix of vaccine response. **A)** Shown are the number of samples for each stimulation of cohort 2. Priority was given for endosomal sensors. All samples could be stimulated the endosomal sensors (TLR3,7,8 9, and cGAS, RIG-1 and STING). B) Correlation heatmap reporting Spearman correlation coefficients (r) as color and number and P values as asterix for each comparison. Statistical differences were assessed with Spearman correlations (B). \**P* < 0.05; \*\**P* < 0.01; \*\*\**P* < 0.001; \*\*\*\**P* < 0.0001.

**Supplementary Figure 10.**
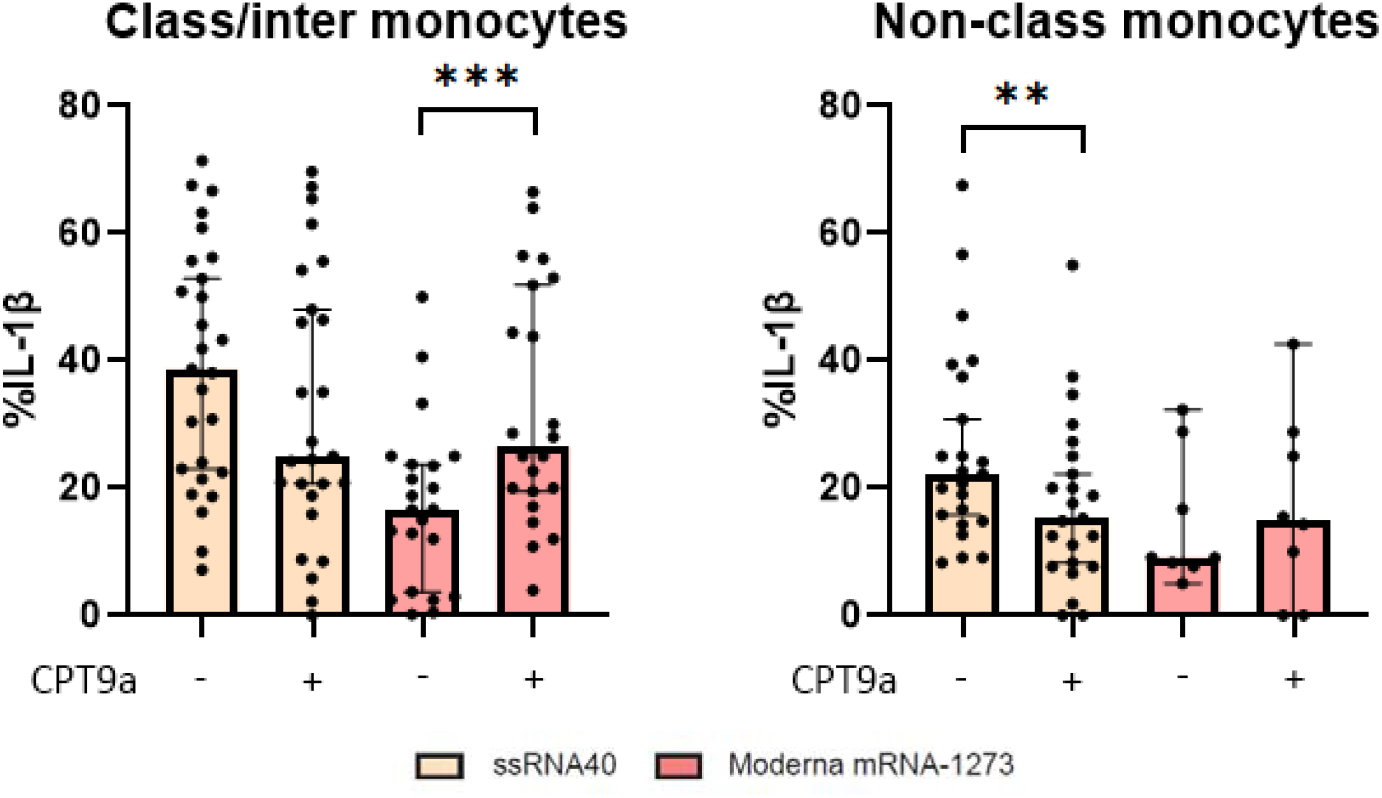
Response of myeloid-derived innate populations to moderna mRNA-1273. PBMCs were stimulated with TLR8 agonist ssRNA40 and Moderna mRNA1273 with and without TLR8 inhibitor CPT9a. Production of IL1β was measured for classical/intermediate monocytes and non-classical monocytes. Statistical differences were assessed with Wilcoxon signed-rank tests. \**P* < 0.05; \*\**P* < 0.01; \*\*\**P* < 0.001; \*\*\*\**P* < 0.0001. Bar plots depict median with 95%CI

## Supplementary tables

**Supplementary Table 1.** Panel of different monoclonal antibodies used for analysis of the TLR expression of immune cell populations.

| # | Marker | Fluorochrome | Company | Cat number | Clone | Dilution | ref control |
| --- | --- | --- | --- | --- | --- | --- | --- |
| 1 | TLR4 | BV421 | Biolegend | 312811 | HTA125 | 1/25 | beads |
| 2 | TLR2 | Pe-Cy7 | Biolegend | 309721 | TL2.1 | 1/50 | beads |
| 3 | CD11c | BUV615 | BD Biosciences | 752323 | 3.9 | 1/50 | cells |
| 4 | CD14 | BV570 | Biolegend | 301831 | M5E2 | 1/100 | beads |
| 5 | CD303 | BUV661 | BD Biosciences | 749920 | V24-785 | 1/100 | beads |
| 6 | TCRgd | PerCP-Vio 700 | Miltenyi Biotec | 130-114-040 | REA591 | 1/120 | cells |
| 7 | CD56 | BUV737 | BD Biosciences | 612767 | NCAM16.2 | 1/200 | cells |
| 8 | CD7 | BV480 | BD Biosciences | 566161 | M-T701 | 1/200 | beads |
| 9 | CD1c | BV711 | Biolegend | 331535 | L161 | 1/200 | cells |
| 10 | CD141 | BV750 | BD Biosciences | 747244 | 1A4 | 1/200 | cells |
| 11 | CD45 | AF700 | Biolegend | 368514 | 2D1 | 1/200 | cells |
| 12 | CD3 | PE-Cy5 | BD Biosciences | 561007 | UCHT1 | 1/400 | cells |
| 13 | HLA-DR | BV650 | Biolegend | 307649 | L243 | 1/400 | cells |
| 14 | CD16 | BUV563 | BD Biosciences | 748851 | 3G8 | 1/800 | cells |
| 15 | CD8 | Spark Blue 550 | Biolegend | 344759 | SK1 | 1/800 | beads |
| 16 | CD4 | cFLuor BYG750 | CYTEK | SKU R7-20160 | SK3 | 1/2000 | cells |
| 17 | TLR3 | APC | R&D systems | IC1487A | poly | 1/50 | beads |
| 18 | TLR7 | FITC | Biolegend | 376907 | S18024F | 1/50 | beads |
| 19 | TLR8 | PE | Biolegend | 395503 | S16018A | 1/50 | beads |
| 20 | TLR9 | apc-f810 | Biolegend | 394811 | S16013D | 1/50 | beads |

